# Time-resolved scanning ion conductance microscopy for three-dimensional tracking of nanoscale cell surface dynamics

**DOI:** 10.1101/2021.05.13.444009

**Authors:** Samuel M. Leitao, Barney Drake, Katarina Pinjusic, Xavier Pierrat, Vytautas Navikas, Adrian P. Nievergelt, Charlène Brillard, Denis Djekic, Aleksandra Radenovic, Alex Persat, Daniel B. Constam, Jens Anders, Georg E. Fantner

## Abstract

Nanocharacterization plays a vital role in understanding the complex nanoscale organization of cells and organelles. Understanding cellular function requires high-resolution information about how the cellular structures evolve over time. A number of techniques exist to resolve static nanoscale structure of cells in great detail (super-resolution optical microscopy^1^, EM^2^, AFM^3^). However, time-resolved imaging techniques tend to either have lower resolution, are limited to small areas, or cause damage to the cells thereby preventing long-term time-lapse studies. Scanning probe microscopy methods such as atomic force microscopy (AFM) combine high-resolution imaging with the ability to image living cells in physiological conditions. The mechanical contact between the tip and the sample, however, deforms the cell surface, disturbs the native state, and prohibits long-term time-lapse imaging. Here, we develop a scanning ion conductance microscope (SICM) for high-speed and long-term nanoscale imaging. By utilizing recent advances in nanopositioning^4^, nanopore fabrication^5^, microelectronics^6^, and controls engineering^7^ we developed a microscopy method that can resolve spatiotemporally diverse three-dimenional processes on the cell membrane at sub-5nm axial resolution. We tracked dynamic changes in live cell morphology with nanometer details and temporal ranges of sub-second to days, imagining diverse processes ranging from endocytosis, micropinocytosis, and mitosis, to bacterial infection and cell differentiation in cancer cells. This technique enables a detailed look at membrane events and may offer insights into cell-cell interactions for infection, immunology, and cancer research.

---

Visualizing dynamic structural changes in live eukaryotic cells at the nanoscale is essential to understand the mechanisms by which cellular components fulfill their function in key cellular processes. The time frame of those processes ranges from seconds/minutes for events such as endocytosis, to hours/days for cell differentiation. To study long-term biological mechanisms, light-based time-lapse microscopy (TLM) methods have been implemented, capturing a sequence of images at regular intervals^8,9^. Among the plethora of TLM techniques on eukaryotic cells, phase-contrast, differential interference contrast and fluorescence microscopy are the most relevant. Visualizing morphological changes in the cell membrane, however, is challenging as they involve dynamic nanostructures with dimensions below the diffraction limit of light^10,11^. Super-resolution microscopy methods have become invaluable in studying live cells^12^. Yet, due to the axial resolution, photobleaching, and phototoxic effects it remains challenging to resolve the membrane surface of a living cell using light-based TLM techniques. In particular, determining the exact boundary of the cell membrane is often impossible at the nanometer scale^13^. Atomic force microscopy offers excellent axial resolution and can be used to image live cells. However, the forces applied by the cantilever tip deform the cell, thereby revealing the cytoskeleton, rather than the native shape of the cell membrane^14^. In addition, the tip-sample forces deform soft structures such as microvilli, making them difficult to resolve^15^. Maintaining long imaging times with AFM on mammalian cells is technologically very difficult due to the complexity and time variance of the tip-sample force interaction. This drastically decreases the viability of cells, in particular those of mammalian origin. High-speed AFM imaging is limited to relatively small imaging ranges, and the need for small AFM cantilevers limits the height of the samples that can be imaged. The ultimate method for single-cell biology would combine the high-resolution 3D information of scanning probe microscopy, with the imaging range, the long-time time-lapse capability, and the high temporal resolution of optical microscopy.

Scanning Ion Conductance Microscopy (SICM)^16^ has been shown to be particularly well suited for imaging soft samples^15,17,18^. By measuring the distance-dependent current through a nanopipette, SICM yields high-resolution images of fragile membrane features, without touching the cell. Possibly the most notable evolution in SICM was the development of backstep/hopping mode by Happel^19^ and the seminal work by Novak and Korchev^18^, making SICM an ideal starting point for long time-lapse imaging of mammalian cells. However, hopping mode SICM is traditionally a rather slow technique with acquisition times in the order of tens of minutes per image, which prevents the study of fast dynamic events.

The imaging speed in SICM is largely determined by the hopping rate^18^ and pipette velocity at which the pipette approaches the surface and retracts when the drop in ion current reaches a certain set point threshold (Fig. 1a in red). Usually, a set point below 2% of the baseline current is used to avoid pipette/sample crashing. The hopping rate is limited by the speed at which the piezo scanner can retract the pipette without leading to excessive overshoot. Once the current drop reaches the set point, the SICM controller tries to retract the pipette as fast as possible. This motion creates an inertia that excites the resonance frequency of the piezo actuator. The larger the pipette velocity is at the time the current reaches the set point, the stronger is this parasitic excitation of the scanner resonances. The resonances are also excited stronger the closer the hopping rate is to the resonance frequency of the piezo actuator. To overcome the second issue, previous efforts have been made in designing SICM scanners with high resonance frequencies which unfortunately goes at the expense of the Z-range^20,21^. Imaging of Eukaryotic cells, however, requires piezo actuators with long-range (>10-20 μm). This traditionally leads to a tradeoff between resonance frequency and range of the actuator. In addition to using a custom high-bandwidth, large-range actuator (Fig. 1b,c,d and Supplementary Fig. 1,2), we overcame this tradeoff in our instrument on the one hand by adaptively slowing down the pipette velocity already before it reaches the trigger setpoint^22^. We use an adaptive gain applied to the piezo motion as a function of the current interaction curve (Supplementary Fig. 2a,b). In addition, the motion dynamics of the actuator are re-shaped by a data-driven controller^7^, implemented in a 16th-order discrete-time filter that compensates the resonances of the electromechanical actuators (Supplementary Fig. 2c-e and Supporting Notes 1). The adaptive hopping mode and the data driven controller increased the achievable hopping rate by a factor of 5 (Fig. 1c,d).

**Fig. 1.**
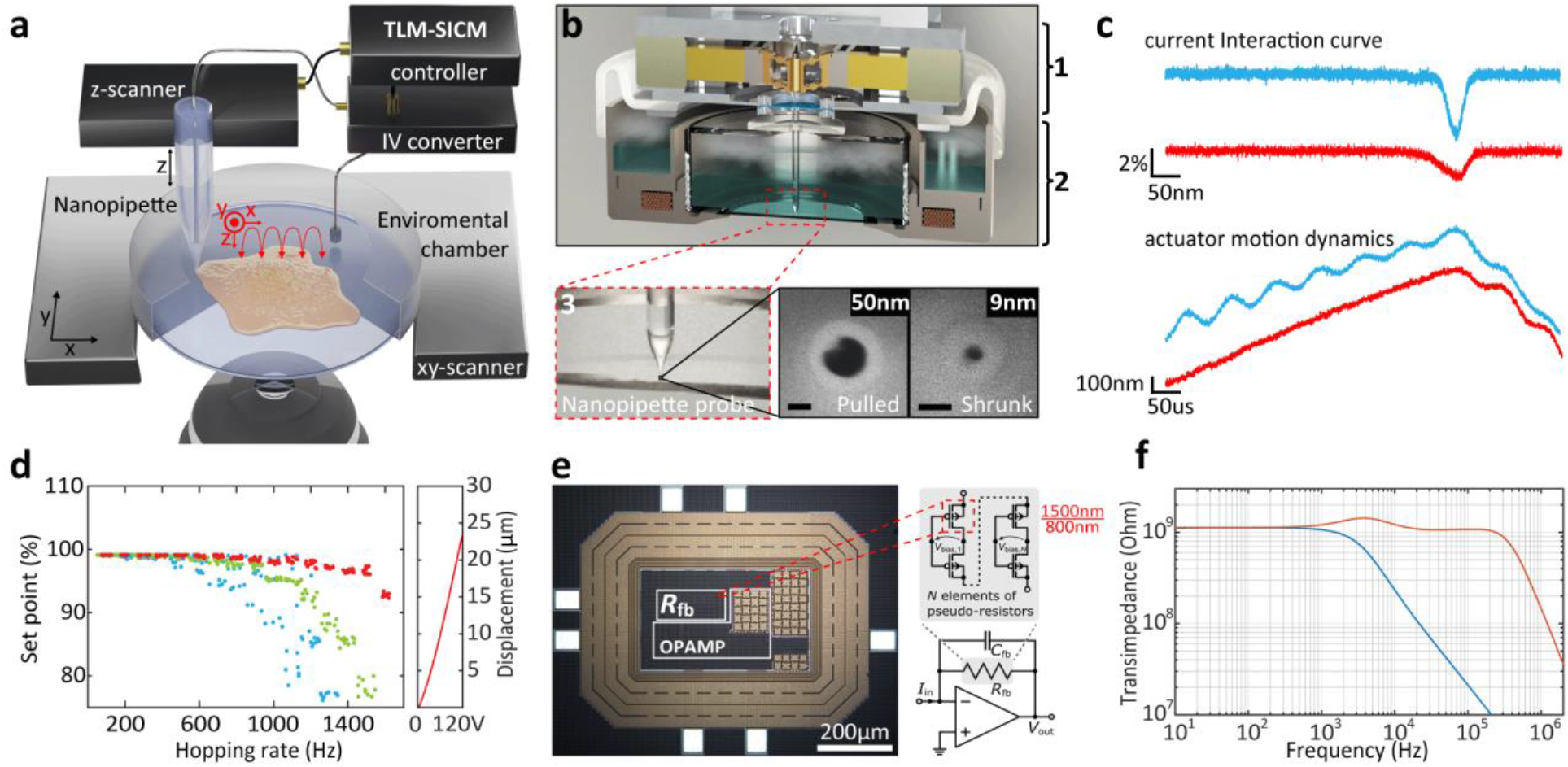
Time-resolved SICM principle and implementation. **a**, Illustration of the SICM principle. **b**, Schematic rendering of a cross-section of the time-resolved SICM system based on a high-bandwidth large-range SICM actuator (1), integrated into a miniature incubator (2). Nanopipettes are fabricated through laser pulling (50nm radius); and successively shrunk through SEM radiation to sub-10 nm pore radius, SEM images (3). Scale bars, 50 nm. **c**, SICM interaction curve (top), and actuator motion dynamics (bottom) with conventional hopping mode in blue and with the time-resolved SICM implementation (adaptive hopping mode and data-driven controller) in red. 1 kHz hopping rate (1.25 mm s^-1^ approach velocity), 1 μm hopping height and 98% set point. **d**, Actuator hopping rate performance with the conventional hopping mode (blue), adaptive hopping mode (green), and adaptive hopping mode together with data-driven controller (red). 1 μm hopping height and 99% set point. **e**, Die micrograph of custom TIA composed of a low-noise operational amplifier and a pseudo-resistor in its feedback. The pseudo-resistor consists of N series-connected pMOS transistor pairs with specific biasing to achieve a large and precise resistance value. **f**, Transimpedance measurements of the 1^st^ stage TIA (blue) and the overall transimpedance using the subsequent amplifier stage for bandwidth extension (red).

Another essential aspect for fast and accurate feedback response is the bandwidth of the low noise current to voltage conversion. Rosenstein et al. have demonstrated that increased performance in nanopore sensing platforms is obtainable by integrating custom tailored complementary metal oxide semiconductor (CMOS) amplifiers^23^. We developed a custom, bandwidth-extended transimpedance amplifier (BE-TIA), which was manufactured in a 180-nm CMOS silicon-on-insulator technology for decreased parasitic well capacitances and leakage currents. It has a subsequent discrete amplifier stage to further increase the overall bandwidth^6^. The TIA’s feedback resistor is set to a very large value of 1 GΩ to achieve low noise and is composed of a multi-element pseudo-resistor (Fig. 1e), i.e., a large number of series-connected small-sized p-channel MOS transistors of W/L = 1500/800 nm with inherent linearization and a specific biasing circuit that facilitates a precise and tunable high-value resistance (Fig. 1e, Supplementary Fig. 4 and Supporting Notes 2). The subsequent amplifier increases the overall bandwidth from 10 kHz to 600 kHz (Fig. 1f). The BE-TIA multiplied the achievable hopping rate in our time resolved SICM by a factor of 8 (Supplementary Fig. 5).

Maintaining cell viability during long-term time lapse imaging requires accurate control of temperature, humidity and CO2 levels in the sample area, without adversely affecting the SICM performance. We integrated our TL-SICM with a custom miniature incubator (Fig. 2a and Supplementary Fig. 6), which ensured cell viability for well over 48h (fig. 2b). Fig 2c and Supplementary Movie 1 shows a 28 hours continuous time-lapse imaging sequence of kidney cells revealing highly dynamic protrusions on the apical cell surface (Fig. 2c) such as lamellipodia (“LP” in Fig. 2c-1,2), and ruffles (“R” in Fig. 2c-1,3), which play a major role in cell motility (Supplementary Fig. 7). Furthermore, we observed filopodia, which are finger-like protrusions that cells use for probing the environment (“FL” in Fig. 2c-1, and Supplementary Fig. 7). We also identified stress fibers that have an essential role in cell adhesion, migration, and mechanotransduction (“SF” in Fig. 2c-1,3). Additionally, we observed cell-to-cell contacts that play a fundamental role in cell-communication and the development of multicellular organisms (“CC” in Fig. 2c-1,2,4,5, and Supplementary Fig. 7). Figures 2d-4,5, and Supplementary Fig. 8 reveal how the interfaces of two cells fuse in syncytium. Of particular interest is the appearance and disappearance of circular dorsal ruffles (“CDR1” and “CDR2” in Fig. 2c-5 to 10, and Supplementary Fig. 9,10). Being able to observe the dynamics of these diverse structures within one long experiment lets us correlate the seemingly independent structures and investigate possible interdependencies.

**Fig. 2.**
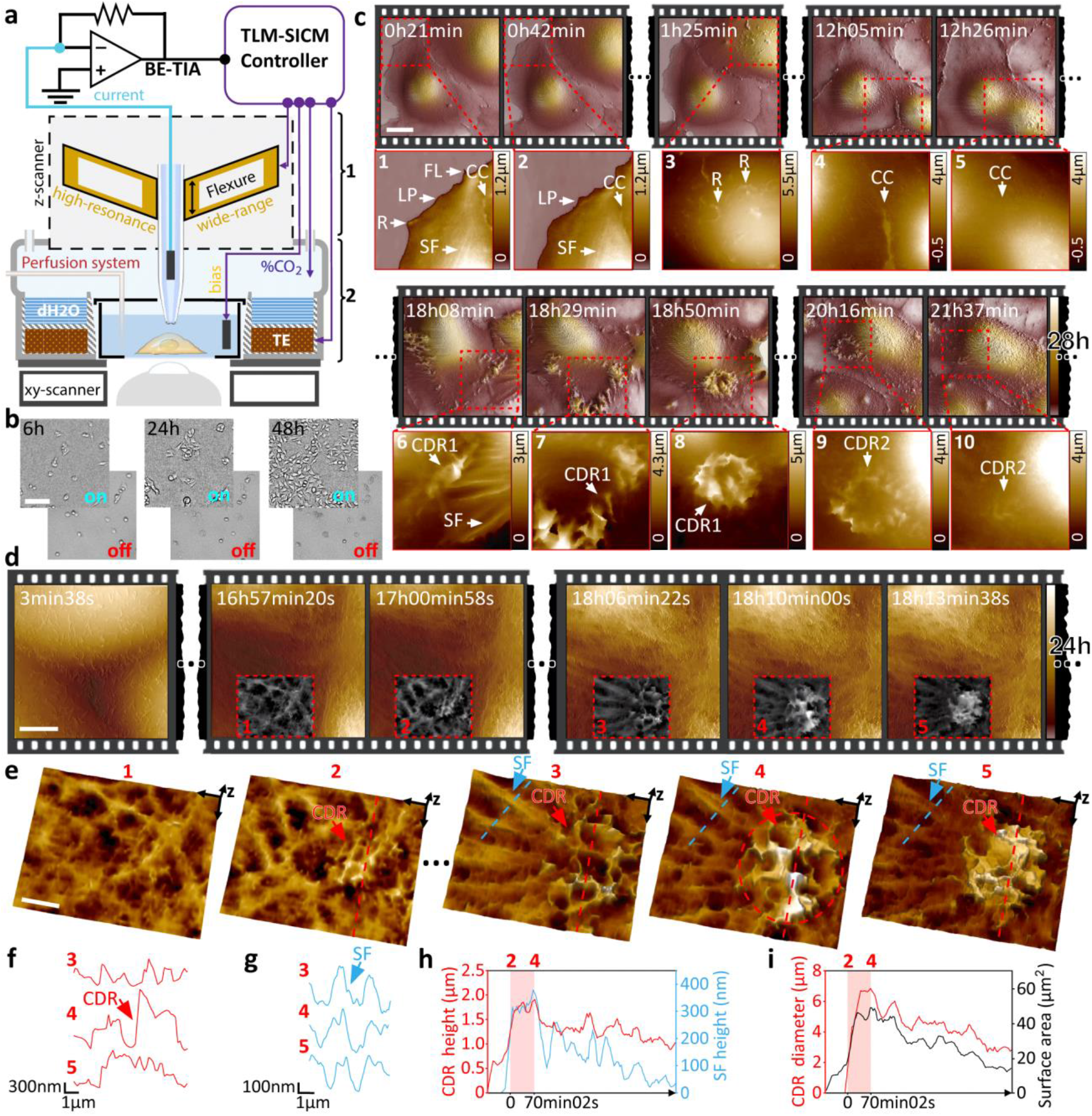
Time-resolved allows long-term 3D visualization of the eukaryotic cell membrane and the tracking of dynamic structures at nanometer resolution. **a**, High-performance pipette z-actuator (1) is integrated into a miniature incubator (2). **b**, Demonstration of cell viability and proliferation over 48 hours in our time-resolved SICM system, with controlled atmosphere ON vs OFF. **c**, 28 hours time-lapse scanning of live kidney cells (COS-7) in DMEM-Hi glucose medium. Scale bar, 20 μm. Several membrane structures can be visualized: ruffles (R), lamellipodia (LP), filopodia (FL), cell-cell contacts (CC), stress fibers (SF), and CDR. **d**, 24 hours time-lapse scanning of live kidney cells tracking the formation of a CDR structure. Scale bar, 5 μm. **e**, 3D view of the CDR with the Initiation spot in panels 1-2, and maximum expansion in panels 3-5. Scale bar, 2 μm. **f**, Height-profile of the CDR ring (red arrow in F). **g**, Height-profile of a nearby SF protruding the cell surface (blue arrow in F). **h**, Plot of CDR and SF max-height, showing a strong correlation during the expansion period (2-4); **i**, Plot of CDR ring diameter and surface area; over 130 frames.

Circular dorsal ruffles (CDR) are elusive dynamic structures, likely related to cell migration, macropinocytosis, and internalization of membrane receptors^24^. These important roles are receiving increasing attention in the scientific community due to their involvement in cancer progression and the facilitation of the pathogen infection^25^. While most studies report that CDRs appear in cells stimulated with mitogenic and/or motogenic factors, our long time-lapse measurements on kidney cells show that these transient structures can appear in optimal culture conditions without external stimulation (Fig. 1d-6 to 10, and Fig. 1e). With time-resolved SICM we tracked a CDR that appeared at 17h00min of scanning with the maximum expansion of the wave reached at 18h10min (Fig. 2,e,f, and Supplementary Movie 2). Recently, Bernitt et al have modeled the appearance of CDRs and associated them with actin polymerization^26^ using confocal microscopy. Our time-resolved SICM data corroborate their findings, adding three-dimensional details of the process at sub-100 nm lateral and sub-10 nm axial resolution. Fig. 1f clearly shows the proximity of the ring-shaped actin mediated wave (CDR) (Fig. 1g) to stress fibers (SF) (Fig. 1h). The polymerization of the stress fibers extrudes the cell membrane and correlates with the formation of the CDR ring (Fig. 1i,j). After expansion, these fibers gradually fade from the surface topography whilst CDR decreases in height and diameter, suggesting their interdependence.

Many of the processes on the level of whole cells occur at a time scale of minutes or hours and can therefore be tracked easily with time-lapse SICM. Sub-cellular events such as endocytosis or infection, however, often occur much faster. Increasing the imaging speed of SICM has thus far been at the expense of imaging volume^15^ to only 7’000μm^3^. This makes studying natural processes on live cells impractical. Our technique combines the ability to address large imaging volumes up to 220’000μm^3^ at moderately fast speeds with high-speed SICM imaging of small details on live cells (Fig. 3a and Supplementary Fig. 11-13, and Supplementary Movie 3). The small-scale, high-speed data can thus be put in the context of the overall cell morphology and growth patterns. Fig. 3a shows a large area 80 × 80 μm overview scan of a kidney cell. Subsequent zooming-in increased the temporal resolution to frame periods of 0.5 s/image at hopping rates of 2.5 kHz. We tracked protrusions at the cell periphery and on top of the cell membrane (Fig. 3a,b,c; and Supplementary Movie 4). Figure 2d,e show endocytic pit formation on live kidney cells (Supplementary Movie 4). We also observed endocytosis on modified melanoma cells (SKMEL) that co-express clathrin-RFP and dynamin-GFP (Fig. 3f). Figure 3g depicts larger area scans (3 μm field of view) with higher resolution (100 × 100 pixels, Supplementary Movie 5) on the cell membrane to identify endocytic pits, and to extract dynamic key parameters such as pit depth and pit area and plot them as a function of time (Fig. 3h). Another endocytic event imaged at 0.5 s/image, but smaller size scale and lower number of pixels is shown in Figure 3i,j. The large dynamic range of the measurements in terms of scan size (500 × 500 nm to 100 × 100 μm), imaging speed (0.5 s/image to 20 min/image), number of pixels per image (1 kilopixel to 1 Megapixel), and a depth of view (22 μm with axial resolution below 10 nm) greatly enhances the type of biological questions that can be studied with time-resolved SICM.

**Fig. 3.**
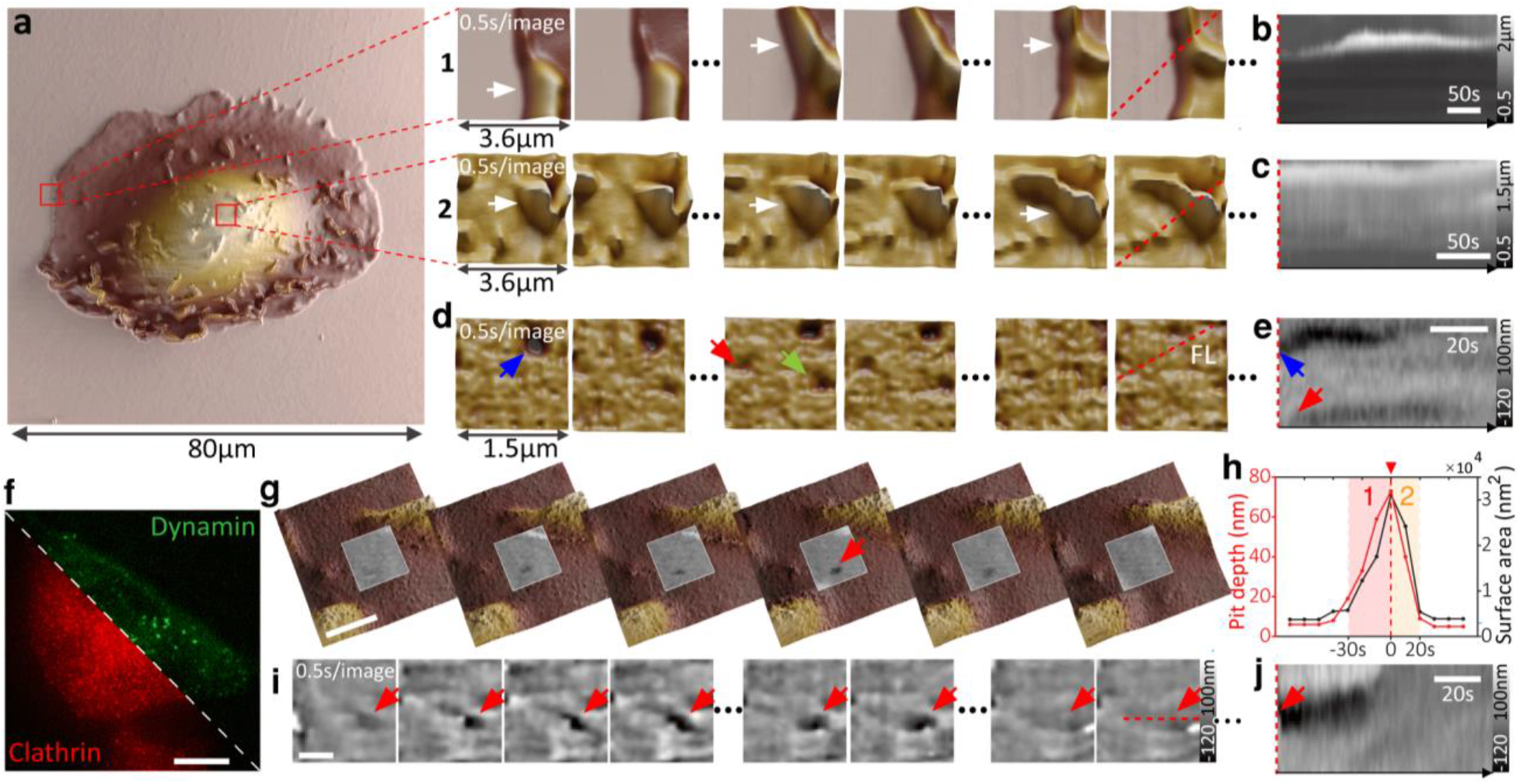
Time-resolved SICM allows a large dynamic scan range essential for long-term monitoring of cells and high-speed performance to track transient biological events at the nanoscale. **a**, Large area scanning of a single kidney cell (80 μm: 512×512 pixels). Fast image acquisition at 0.5 s/image (2.5 kHz hopping rate) on the cell periphery (1), and on top of the cell (2). Arrows point to dynamic ruffles. **b** and **c**, Kymogram showing the dynamics of ruffles over time (red dashed lines in a, 1 and 2). **d**, Fast image acquisition of 0.5 s/image of the kidney cell membrane with arrows pointing to several endocytic events; with the respective kymogram in **e**. **f**, Fluorescence image of a transformed melanoma cell that co-expresses clathrin-RFP and dynamin-GFP. Scale bar, 20 μm. **g**, Fast image acquisition of a larger area (3 μm: 100×100 pixels) at 10 s/image (1 kHz hopping rate) detecting the formation of an endocytic pit (red arrow) within 50 s over 5 data points. Scale bar, 1 μm. **h**, Plot of endocytic pit depth and surface area. 1: growing; and 2: closing. (**i**) Fast image acquisition at 0.5 s/image of an endocytic pit (red arrow) within 50 seconds and 100 data points a 15 nm radius pipette. Scale bar, 500 nm; with the respective kymogram in **j**.

Time-resolved SICM is particularly well suited for biological questions that relate either to a change in 3D cell morphology, or where membrane trafficking plays a major role. Morphological changes of transformed cells, for example, are essential in cancer diagnosis and treatment decision-making^27^, but are difficult to track over long durations. Knowledge about associated events on cell membranes, their dynamics and roles in tumor evolution is limited by a lack of tools for long-term imaging. In cultured melanoma cells (B16-F1), important morphological changes linked to cell differentiation and motility are induced by 3’-5’-cyclic adenosine monophosphate (cAMP) signaling, which is implicated in mediating resistance to BRAF inhibitor therapy^28^, and which can be activated in vitro by treatment with the adenylate cyclase (ADCY) agonist forskolin (FSK)^29^ (Supplementary Fig. 14). To evaluate the spatiotemporal resolution of time-resolved SICM to reveal cAMP-induced changes, we monitored melanoma cell morphology during 48 hours of sustained FSK treatment^30^ (Fig. 4a and Supplementary Movie 6). Within this time frame, FSK treatment drastically increased the outgrowth and branching of interwoven cell dendrites that were frequently suspended >5 μm above the substrate (Fig. 4b, c).

**Fig. 4.**
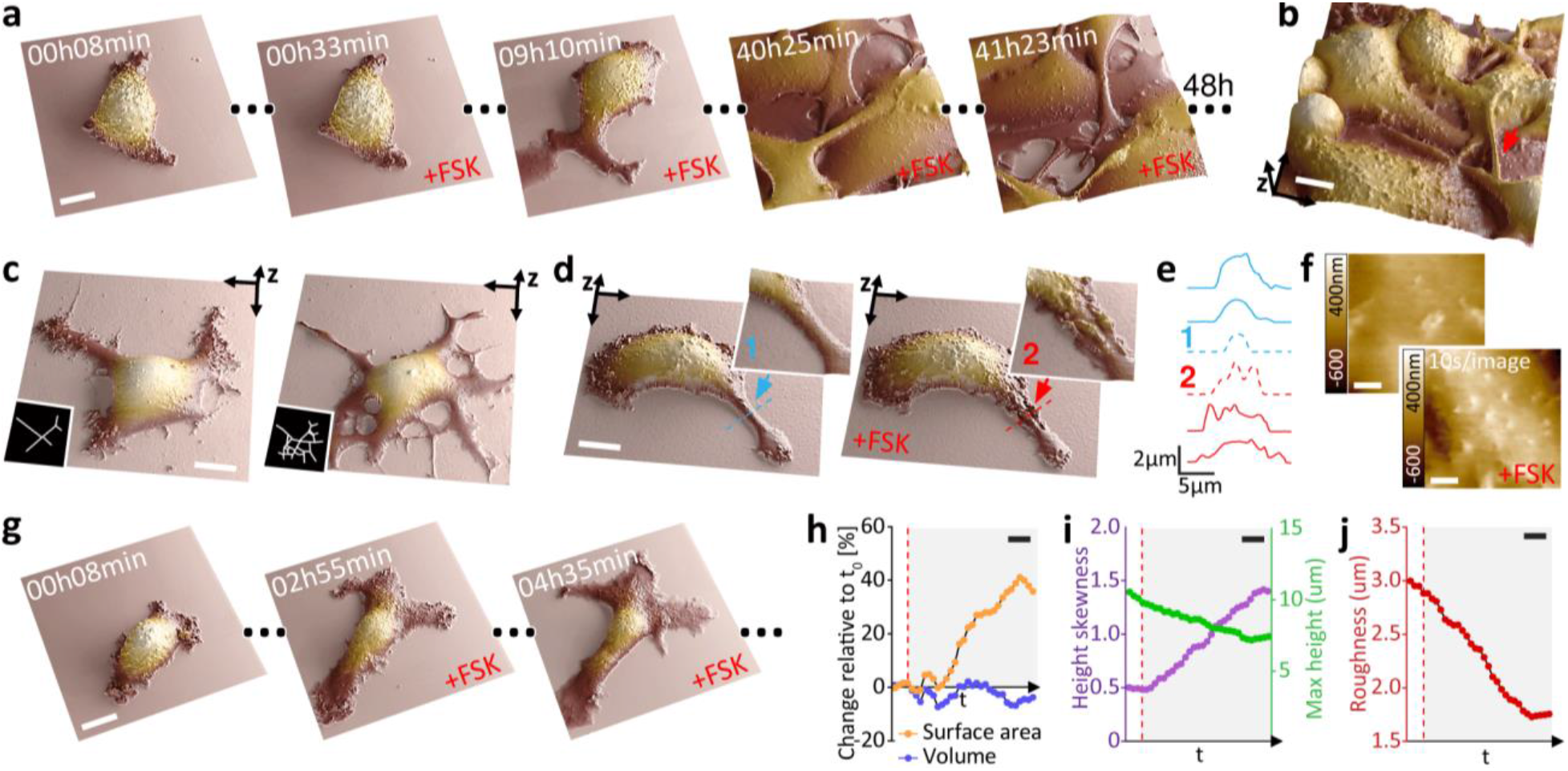
Time-resolved SICM enables a large dynamic scan range of cells and suspended structures, allowing long-term visualization of differentiation and morphological changes in melanoma cells. **a**, 48 hours time-lapse scanning of melanoma cell (B16-F1) differentiation during prolonged treatment with 20μM forskolin (FSK). Scale bar, 10 μm. **b**, The long actuation range allows the nanoscale visualization of dendrites (branched cytoplasmic protrusions), suspended 5 μm above the substrate (red arrow). Scale bar, 10 μm. **c**, FSK-induced dendrite outgrowth. Scale bar, 10 μm. **d**, Tracking a single dendrite before (blue arrow) and after adding FSK (red arrow). Scale bar, 10 μm. **e**, Height profile of the dendrite over a time sequence with 1 and 2 shown in panel d. **f**, Visualization of the real-time effect of FSK on the cell membrane with fast image acquisition at 10 s/image (1 kHz hopping rate). Scale bar, 1 μm. **g**, Visualization of long-term morphological changes associated with FSK-induced melanoma cells differentiation. Scale bar, 20 μm. **h**, Surface area and volume percentage change relative to the first frame. **i**, Maximal height and height skewness. **j**, Membrane roughness. Scale bar, 100 minutes. The dashed line in red represents the addition of FSK.

The high spatial resolution afforded by time-resolved SICM imaging revealed that FSK treatment rapidly increased the width of dendrites (Fig. 4d,e, and Supplementary Fig. 15,16), and revealed the presence of membrane protrusions already during the earliest time points examined (Fig. 4f, and Supplementary Fig. 17, and Supplementary Movie 7). Prolonged FSK treatment resulted in a gradual increase in cell surface area with no significant change in cell volume (Fig. 4g,h, and Supplementary Fig. 18a,b, and Supplementary Movie 8). Curiously, these changes were accompanied by a rise in cell height skewness (Supplementary Fig. 18c), decreased cell height (Fig. 4i and Supplementary Fig. 18d), and by a long-term decrease in membrane roughness (Fig. 4j and Supplementary Fig. 18e). The roughness measured is associated with the presence of membrane protrusions, a feature related to focal adhesions that are mediated by rearrangement in actin cytoskeleton^31^ and increased invasiveness in cancer^32^.

We also applied time-resolved SICM to investigate the mechanisms of bacterial adhesion of mammalian host cells, a first step towards infection by many pathogens. We monitored the binding of E.coli to epithelial cells through the display of synthetic adhesins (on bacteria) that mimic a pathogenic context. These bacterial cells display anti-GFP nanobodies (VHH) and bind to GFP anchored to the plasma membrane of HeLa cells^33^. Interestingly, we observed an increase in the GFP signal around bacteria suggesting a local accumulation of the VHH-GFP bonds that represents an increase in the overall affinity of the bacterium to the host cell membrane over time (Fig. 5,a,b,c and Supplementary Fig. 19). In addition, time-resolved SICM revealed bacteria dividing directly on the host cell membrane, proving that the attachment to the mammalian cell does not prevent bacterial proliferation (Fig. 5d, Supplementary Fig. 20 and Supplementary Movie 9). Occasionally, the binding of the bacterium tightened to the point it triggered its internalization by the mammalian cell (Fig. 5e, Supplementary Fig. 21), as confirmed also by confocal microscopy (Supplementary Fig. 22).

**Fig. 5.**
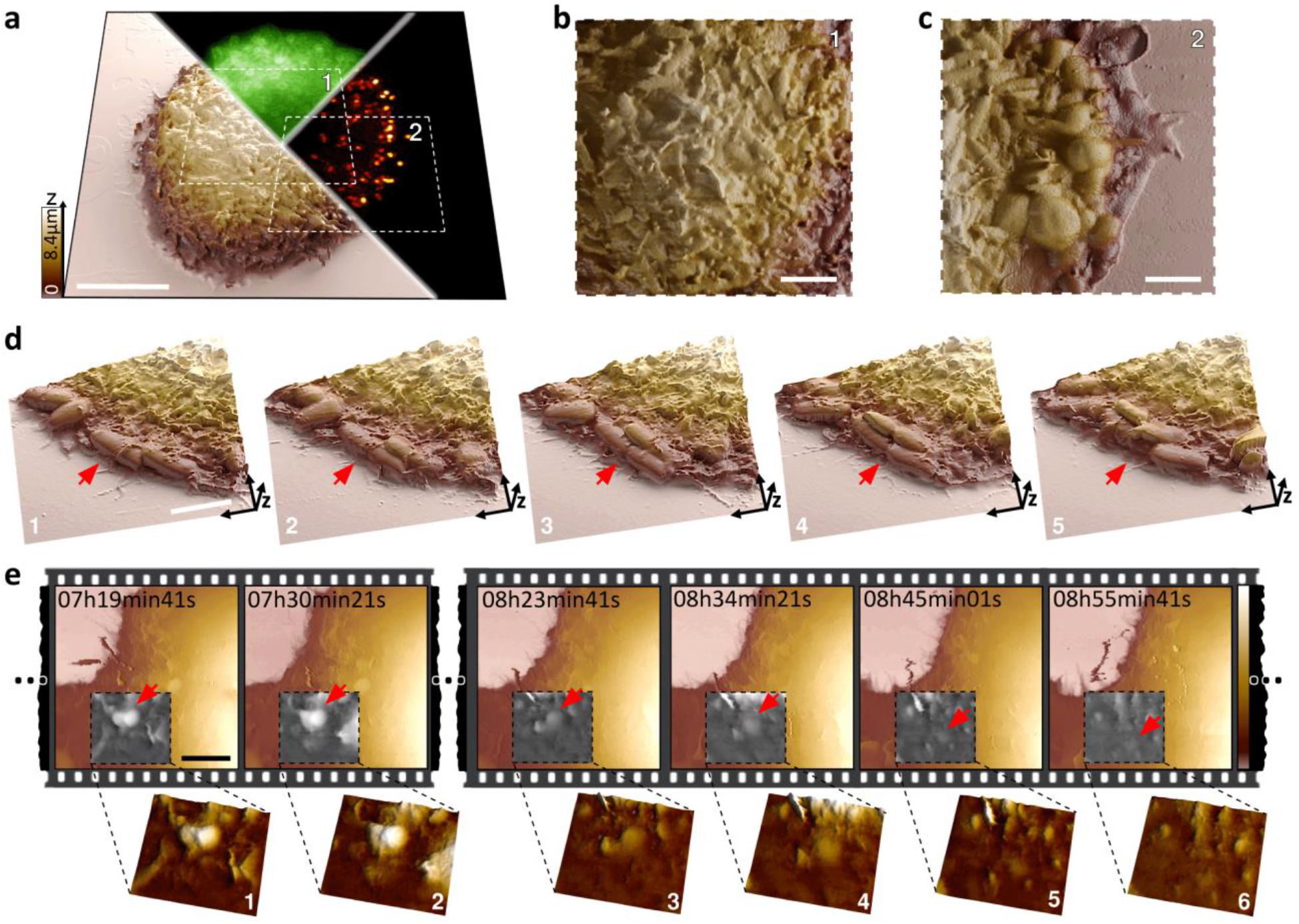
Long-term visualization of infection mechanisms in bacterial adhesion and entry into host cells using time-resolved SICM. **a**, Correlated three-dimensional representation of the live cell membrane surface with SICM. HeLa cells displaying a synthetic GFP receptor enabling its direct visualization (GFP, green channel) sequestered by its ligand: VHH expressed by E. coli (red channel). Scale bar, 10 μm. **b**, E. coli can be resolved on top of the host cell body (area 1 marked in a); **c**, and on the host cell edge (area 2 marked in a). Scale bar, 4 μm. **d**, 3D visualization of bacteria infecting a mammalian cell. The red arrow points to a bacterium adhered to the cell, elongating (1-2) and dividing (3-5) on the mammalian cell membrane. Scale bar, 4 μm. **e**, Live monitoring of bacteria engulfment and internalization; the red arrow points to a bacterium adhered to the host (1, 2) and being internalized (3-6). Scale bar, 4 μm.

Complementary to fluorescence microscopy, time-resolved SICM reveals the precise topography and position of the bacteria relative to the membrane, as well as the pathogen-membrane interactions. SICM is label-free and does not suffer from phototoxicity and photobleaching that could prevent the long-term visualization of these processes with fluorescence microscopy. In contrast to AFM, SICM is a truly non-contact measurement that prevents detachment of the settled bacteria or artificially perturbs their weak interactions with the host cell body that govern infection. SICM imaging did not prevent adhesion and invasion and was able to monitor the infection process over long imaging times. Due to the non-contact nature and minimized perturbations, we will be able to investigate a broad range of infection mechanisms and internalization processes in the future.

We have demonstrated in this work some of the benefits of time-resolved SICM for studying nanoscale processes in cell biology. However, some technological hurdles still need to be overcome for time-resolved SICM to become a routine tool in cell biology. One key requirement for good quality SICM imaging is a high-quality nanopipette with an appropriate aperture diameter for the specific application. While high-speed imaging generally benefits from a lower resistance pipette (i.e. a larger opening), high-resolution requires a small opening^21^. Obtaining reproducible and robust capillaries at precise diameters is still a challenge for many groups starting to use SICM. For SICM to reach its full potential, a reliable off-the-shelf source of SICM probes would be beneficial (similar to what is available for AFM cantilevers).

While TL-SICM can obtain unprescedented characterization of the cell shape and surface characteristics, these aspects often need to be correlated with biochemical information and changes to the internal organization of the cells. The combination of SICM with high performance optical microscopy is a very promising area^34^. The TL-SICM is integrated with an inverted optical microscope (Supplementary Fig. 23) and can thus be used with many of the recently developped super resolution microscopy techniques^35^.

The SICM field has developed rapidly in recent years, with many additional measurement capabilities being added, such as sample stiffness^36^, surface charge^37^, and local pH^38^. Combining these developments with time-resolved SICM or high-speed SICM will provide unprecedented insights into eukaryotic membrane processes with three-dimensional nanometer detail.

## Acknowledgments

The authors thank Navid Asmari and Santiago Andany for assistance. We also thank Mustafa Kangül, Matthias Neuenschwander, Mélanie Hannebelle, Benjamin Rothe, Pierpaolo Ginefra and Patrick Frederix for their valuable discussion. We thank Prof. Aurélien Roux, Prof. Hilal Lashuel, Anne-Laure Mellier and Veronika Cencen for providing biological samples. We thank ATPR (EPFL) for mechanical fabrication.

## Author contributions

S.L. conceived the idea, designed and built the instrument, designed experiments, prepared samples, performed measurements, analyzed data, wrote the final manuscript. B.D. conceived the idea, designed and built the instrument, designed experiments, prepared samples, performed measurements and analyzed data. K.P. designed experiments, prepared samples, analyzed data, and together with D.C. gave valuable input on the application in melanoma. X.P. designed experiments, prepared samples, and together with A.P. gave valuable input on the application in infection mechanisms of bacteria. A.P.N. and C.B. developed controller hardware and software. D.D. and J.A. designed and built the TIA chip for high-speed current to voltage conversion. V.N. processed data and together with A.R. gave valuable input on fluorescence correlative data. G.E.F. conceived the idea, designed the instrument, supervised the project and wrote the final manuscript. All authors edited the manuscript.

## Methods

### Cell lines and cell culture

In this study, we used monkey kidney fibroblast-like cells COS-7, SKMEL human melanoma cells, B16-F1 mouse melanoma cells (ATCC), and HeLa ovarian cancer cells. COS-7 and SKMEL cells, modified to co-express dynamin-GFP and clathrin-RFP^39^ were cultured in DMEM high glucose without phenol red medium (Gibco, Thermo Fisher Scientific), containing 10% of fetal bovine serum (Gibco, Thermo Fisher Scientific), 1% penicillin-streptomycin (Gibco, Thermo Fisher Scientific) and 4 mM L-glutamine (Gibco, Thermo Fisher Scientific). B16-F1 mouse melanoma cells were maintained in DMEM (Sigma) supplemented with 10% fetal bovine serum, 50 μg/ml gentamicin (Gibco, Thermo Fisher Scientific), and 1% GlutaMAX (Gibco, Thermo Fisher Scientific). Cells were regularly tested negative for *Mycoplasma* (SouthernBiotech 13100-01). HeLa cells were stably engineered to display a doxycycline-inducible GFP fused to a mouse CD80 transmembrane domain using standard second-generation lentivector production protocols and the plasmids pMD2G (Addgene 12259), pCMVR8.74 (Addgene 22036), and pXP340^33^ [eGFP(N105Y, E185V, Y206F)_mCD80_C-terminal into pRRLSIN.cPPT.GFP.WPRE (Addgene 12252)]. Before co-culture experiments, HeLa cells were seeded at 50,000 cells/ml on 35 mm glass dishes (Ibidi) in 400 μL Fluorobrite supplemented with 10% FBS and 1% Glutamax (Life Technologies). After 4-6 hours of attachment, the medium was renewed and supplemented with 500 ng/ml doxycycline (HiMedia) for the induction of GFP display, and with 3 μL of Deglycosylation Mix II (New England Biolab) to digest the glycocalyx and ease the access of bacteria to the cell membrane. *E. coli* K12 (BW25113), a flagellated (deltaFliCDST) and stably expressing mScarlet were retransformed with pDSG339 for tetracycline-inducible VHH anti-GFP display^40^. Stationary precultures of bacteria were diluted 1:3000 and induced overnight with 250 ng/ml tetracycline (Sigma) in LB.

### Time-resolved SICM instrumentation

The time-resolved SICM setup consists of a custom-built pipette Z-actuator integrated into a controlled-atmosphere device, critical for cell viability during imaging. The temperature is regulated with a temperature controller from THORLABS (TC200), and the percentage and flow rate of CO2 gas infusion is regulated with a gas mixer from Life Imaging Services (The Brick). While keeping optimal cell culture conditions of 37 °C and 5 % CO2 for several days, the high-performance Z-actuator configuration achieves a wide mechanical displacement amplification of 22 μm scanning range on the cell surface; and preserved high-speed SICM performance with the first resonance at above 13 kHz; The Z-actuator is driven by a custom made low-phase lag piezo controller^4^, and integrated with a stepper-motor stage for sample approach. The XY-scanner used was a Piezo-Nanopositioning stage with 100 μm x-y travel range on the sample (P-733 Piezo NanoPositioner, Physik Instrumente), driven by a low-voltage piezo amplifier (E-500 Piezo Controller System, Physik Instrumente). The XY-scanner is assembled in a custom-built micro translation stage, mounted atop an inverted Olympus IX71 microscope body. Therefore the system offers correlative SICM capabilities with fluorescence microscopy^35^. For fluorescence excitation, we used a four-color (405 nm, 488 nm 561 nm, 647 nm) pigtailed Monolithic Laser Combiner (400B, Agilent Technologies), controlled by a custom-written LabVIEW software. Images were acquired with a sCMOS camera (Photometrics, Prime 95B) and Micromanager software. The nanopipettes used as a probe in SICM were made of Borosilicate (Sutter Instrument) and quartz (Hilgenberg GmbH). They were fabricated with a CO2 laser puller (Model P-2000, Sutter Instruments) with a radius from 60 nm to sub-nanometer sizes. Quartz capillaries can be shrunk to a sub-10 nm radius using electron beam irradiation^5,41^. The pipette current was amplified by a transimpedance amplifier NF-SA-605F2 (100MΩ gain, NF corporation) and a 1GΩ custom-made TIA (Supplementary Notes 2) for high-speed current to voltage conversion^6^ (Fig. 3a,g). A low-pass filter setting between 10-100 kHz was implemented after the pre-amplifier (Stanford SR560). A custom SICM controller (Supplementary Notes 1) consisting of a data-driven scan engine^7,42^ and adaptive hopping mode was implemented in LabVIEW on a NI USB-7856R OEM R Series (National Instruments, Austin, TX, USA) in the framework of a home-built extensible high-performance scanning probe controller.

### Time-resolved SICM measurements

The TL-SICM measurements were performed in the optimal culture conditions mentioned above at 37 °C and 5 % CO_2_COS-7 and SKMEL cells were seeded at 30,000 cells/ml in 35 mm glass dishes (Ibidi) for the long time-lapse experiments and 50,000 cells/ml for the high-speed measurements. For transfected SKMEL cells, the surface was coated with poly-D-lysine (Gibco, Thermo Fisher Scientific). Coverslips were washed with PBS before seeding the cells in 2 ml of DMEM medium. Prior to the experiments, the cells were grown overnight (12-16 h) at 37 °C and 5 % CO_2_. In experiments using B16-F1 mouse melanoma cells, 50,000 cells/ml were seeded in 35 mm glass dishes (Ibidi) in 2ml volume of completed culture medium. After overnight culture (13-16 hours), cells were used for scanning. Forskolin (10-2073 Focus Biomolecules) was added to the cells through a perfusion system integrated with the environmental mini-chamber to a final concentration in the cell medium of 20 μM FSK. In the cell infection co-culture measurements, HeLa cell supernatant was renewed to 2 ml medium and 200 μL of stationary bacterial culture was added for 30 minutes (Fig. 5a,b,c) and 15 minutes (Fig. 5d), before co-culture. The samples were intensively washed 5 times with PBS. 2 ml of DMEM high glucose without phenol red medium (Gibco, Thermo Fisher Scientific) containing 10% fetal bovine serum (Gibco, Thermo Fisher Scientific) was added prior to imaging. In the long time-lapse experiments (Fig. 5e), stationary bacteria (100 μL) were added to the medium through a perfusion system integrated into the environmental mini-chamber after 1 hour of scanning. A DC voltage bias in a range of +150 to +300 mV was applied to the pipette. The current setpoint used in the hopping actuation was 99% of the normalized current recorded (98% for high-speed imaging). SICM image sequence in Fig. 2c was acquired at 200 Hz hopping rate and 6 μm hopping height, 512×256 pixels. Image sequence in Fig. 2d was acquired at 200 Hz hopping rate and 3 μm hopping height, 256×128 pixels. Image sequence in Fig. 3a,d,e were acquired at 2.5 kHz hopping rate and 200 nm hopping height, 40×25 pixels. Image sequence in Fig. 3g was acquired at 1 kHz hopping rate and 500 nm hopping height, 100×100 pixels. Image sequence in Fig. 4a,b,g were acquired at 125 Hz hopping rate and 6μm hopping height, 300×200 pixels. Fig. 4c was acquired at 200 Hz hopping rate and 5μm hopping height, 256×256 pixels. Image sequence in Fig. 5 at 125 Hz hopping rate and 5μm hopping height, 512×512 pixels (Fig. 5a,b,c) and 256×256 pixels (Fig. 5d,e).

### Data processing and analysis

SICM images were further processed using Gwyddion^43^. Images were corrected for scanline mismatch with a median of differences row alignment and line artifacts characteristic of SPM were corrected. Afterward, a 2-pixel conservative denoise filter was applied. Images were exported with uniform pixels aspect ratios in 16 bit Portable Network Graphics format. For better visibility, the topography channel is merged with the slope channel to highlight the surface structures in all the data sets shown in panels with a video frame. In SICM images with a greyscale color map the background was flattened with a subtraction of a median level. To generate the three-dimensional shape of cells we used the advanced open-source 3D rendering tool Blender 3D for data visualization. Normalized topographical SICM data was imported as a height map and scaled in the axial direction. The corresponding SICM colormap was used as a color, projected on the topography. Videos were generated in ffmpeg format in Fiji^44^.

