## Supplementary information for "Time-resolved scanning ion conductance microscopy for three-dimensional tracking of nanoscale cell surface dynamics"

[1]Laboratory for Bio- and Nano-Instrumentation, Institute of Bioengineering, School of Engineering, Swiss Federal Institute of Technology Lausanne (EPFL); Lausanne, Switzerland. [2]Laboratory of Developmental and Cancer Cell Biology, Institute for Experimental Cancer Research, School of Life Sciences, Swiss Federal Institute of Technology Lausanne (EPFL); Lausanne, Switzerland. [3]Laboratory of Microbial Mechanics, Institute of Bioengineering and Global Health, School of Life Sciences, Swiss Federal Institute of Technology Lausanne (EPFL); Lausanne, Switzerland. [4]Laboratory of Nanoscale Biology, Institute of Bioengineering, School of Engineering, Swiss Federal Institute of Technology Lausanne (EPFL); Lausanne, Switzerland. [5]Institute of Smart Sensors, Universität Stuttgart; Stuttgart, Germany. *

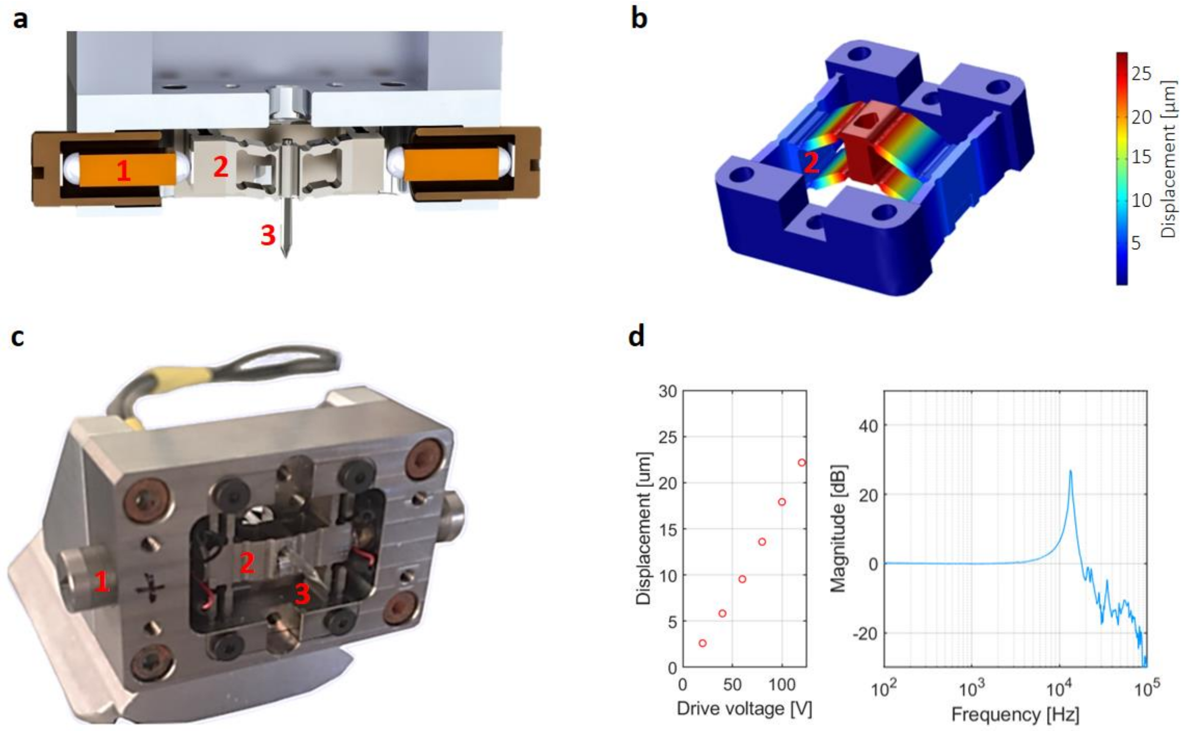


***Supplementary Fig. 1: Long-range actuator with preserved high bandwidth for wide axial scanning range on the cell surface. a**) 3D rendering of a cross-section of the SICM pipette actuator. 1 shows the piezoelectric element, 2 shows the titanium flexure, and 3 the silicate nanopipette. **b**) Finite element simulation of the motion of the actuator configuration allowing for a mechanical amplification with high-resonance. **c**) Implementation of custom-built pipette actuator to perform cell topography imaging. **d**) Mechanical displacement range of 22.17 µm for 120 V drive voltage (on the left in red). **e**) and the actuator frequency response curve with the first resonance at 13.5 kHz (on the right in blue).*

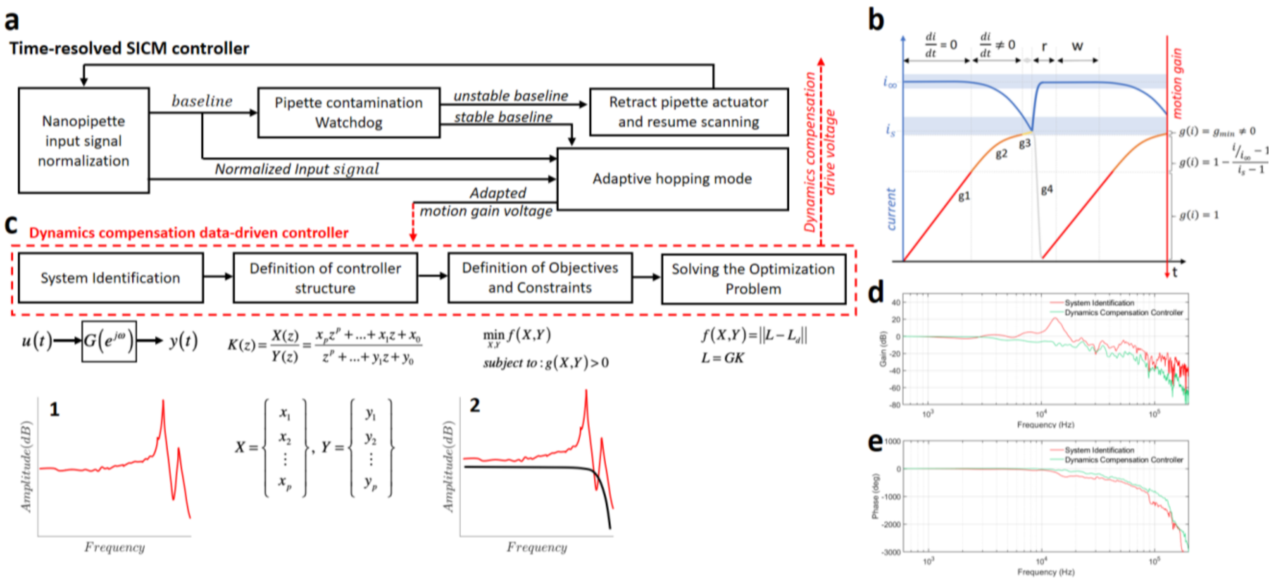


***Supplementary Fig. 2: Working principle of the time-resolved SICM controller. a)*** *Wiring diagram of the controller highlighting the main features: Adaptive hopping mode, a data-driven controller, and a pipette contamination watchdog.* ***b****) The principle of the adaptive hopping mode with an illustration showing the relationship between the current over time (blue) and piezo motion gain (red). The gain (g1 and g2) is proportional to the change in current (di/dt), converging to a minimal gain (g3) until set point is reached, triggering the piezo retraction (g4). The baseline is calculated by recording and averaging data over a time window size (w) after a time delay (r).* ***c****) The data-driven controller[1] is composed of four steps: System identification, Definition of controller structure, Definition of objectives and constraints, and Solving the optimization problem.* ***d****) System identification of the SICM actuator (Red) and its response after the implementation of the data-driven controller (Green).* ***e****) Phase plot showing the system's response before (Red) and after (Green) the data-driven controller implementation.*

### Supplementary note 1: Time-resolved SICM controller implementation

The time-resolved SICM controller consists of three main features (Supplementary Fig. 2a): Adaptive hopping mode, a data-driven controller, and a pipette contamination watchdog. The adaptive mode is characterized by an adaptive gain applied to the piezo motion in the function of the measured current slope. The model used in the adaptive gain is described as $g(i) = 1 - \frac{i/i_\infty - 1}{i_s - 1}$, where $i$ is the measured current, $i_\infty$ is the baseline current (current when the pipette is far from the surface), $i_s$ is the current set point, and $g_{min}$ is the minimum gain (Supplementary Fig. 2b). The adaptive hopping mode is an adaptation of the closed-loop approach-retract-scanning (ARS) model[2].

The data-driven controller design consists of four main steps (Supplementary Fig. 2c). In the first step, a pseudo-random-binary-sequence signal is applied to the input ($u(t)$) of the piezo-actuator and the motion is recorded ($y(t)$). The acquired input-output data is used to construct a non-parametric frequency response ($G(e^{jw})$) of the piezo-actuator which represents its dynamics (c1). Afterward, a structure is defined for the controller based on the available resources. Here, a parametric 16th-order discrete-time filter is chosen, $K(z) = \frac{X(z)}{Y(z)} = \frac{x_p z^p + \dots x_1 z + x_0}{z^p + \dots y_1 z + y_0}$; where $X$ and $Y$ are the parameters that shape the controller dynamics. In order to optimally select these parameters, a set of objectives ($f(X,Y)$) and constraints ($g(X,Y)$) are defined such that the combined behavior of the system (controller and actuator) satisfies our performance expectations. These optimization specifications determine the bandwidth of the system while guaranteeing its stability. For this configuration, the open-loop of the dynamics ($L = GK$) is shaped based on a desired dynamic response ($L_d$). In this regard, the area between the two frequency responses ($\|L - L_d\|_2$) is minimized (c2). A convex optimization method is utilized to design the controller such that it attenuates the high-amplitude dynamics of the piezo-actuator (Supplementary Fig. 2 d,e). This whole design process is executed once and the selected controller parameters are used through the rest of the process to prepare the drive signal of the SICM actuator.

In addition, nanopipette pore contamination is a frequent event in long-term SICM measurements that lead to a decrease in the image quality and often breaks the glass pipette. Therefore, in order to perform long-term imaging a current signal monitoring controller (Pipette contamination watchdog in *Supplementary Fig. 2a*) triggers upon pipette contamination is detected and retracts the pipette from the medium. Capillaries forces successfully remove the contamination and the scanning is resumed, ensuring a long-lasting imaging performance.

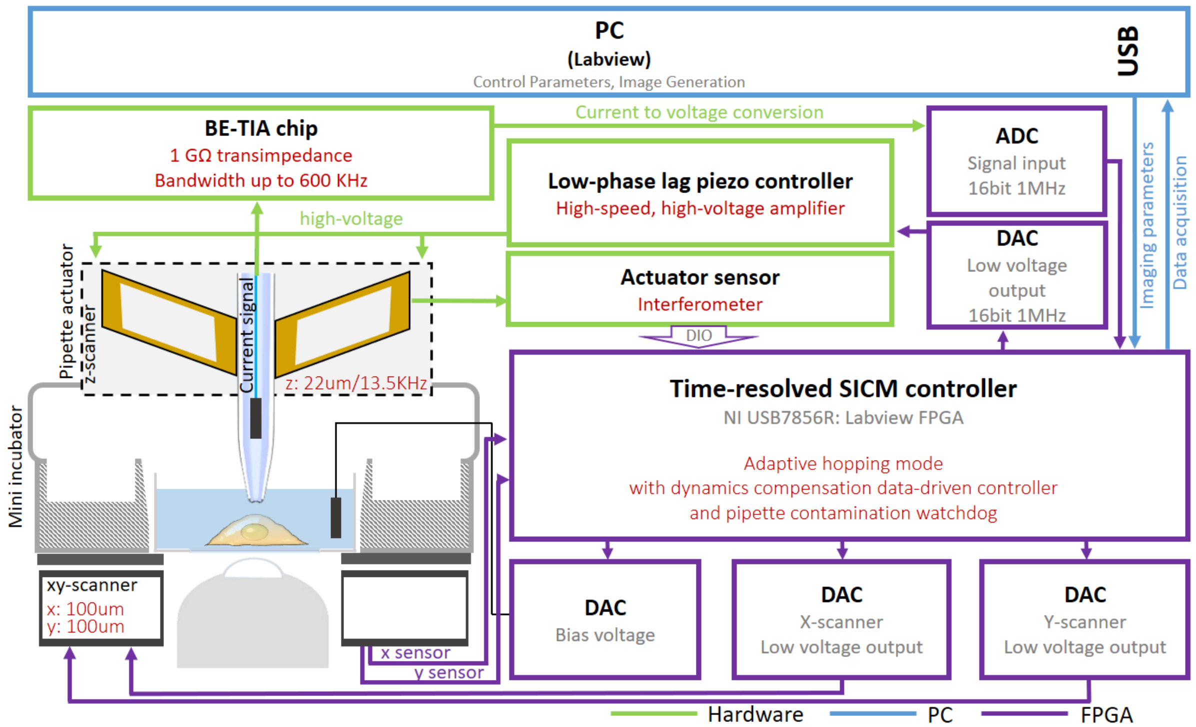


***Supplementary Fig. 3: Schematics of the time-resolved SICM setup implementation.*** *The current generated through the pipette nanopore is converted to voltage (BE-TIA) and amplified with a transimpedance of 1GΩ used as input feedback control signal in the SICM controller (Time-resolved SICM controller). The response of the system is shaped with motion dynamics information from the interferometer (Actuator sensor), such that it attenuates the high-amplitude resonances. The piezoelectric actuators (z-scanner) are driven by a custom made high-voltage amplifier*[3] *with very low-phase lag (Low-phae lag piezo controller), ensuring a fast piezo-actuation response.*

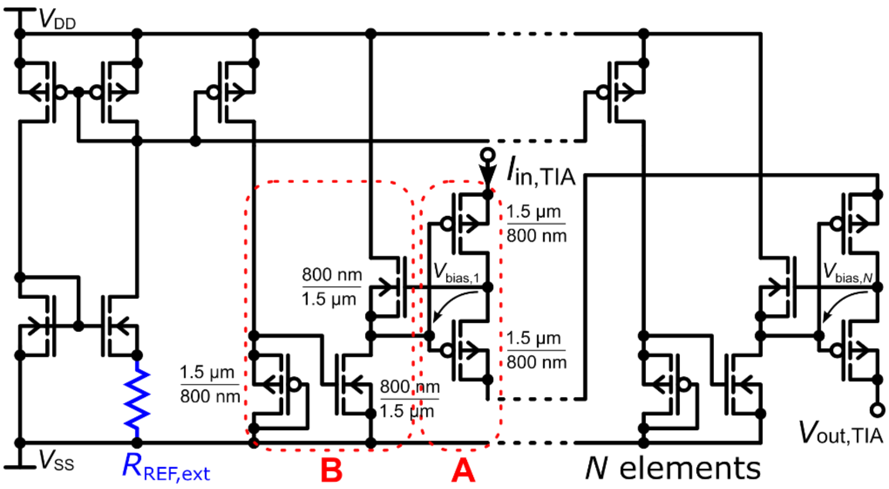


***Supplementary Fig. 4: Schematic of the implemented multi-element pseudo-resistor (MEPR).*** *$I_{in}$ is the TIA current input, $V_{out}$ is the TIA voltage output, and $R_{REF,ext}$ is the external reference resistor, which sets the MEPR's large resistance value. A, p-channel MOS transistor pairs. B, Biasing circuit based on a pseudo current mirror.*

### Supplementary note 2: Multi-element pseudo-resistor (MEPR) implementation

The feedback resistor used in the TIA consists of p-channel MOS transistor pairs, which are biased in weak inversion and operated in the linear regime (A). Such devices are also named as pseudo-resistors due to their resistive behavior for drain-source voltages $V_{DS}$ below the thermodynamic voltage $U_T=kT/q$, of which k is the Boltzmann constant, T the absolute temperature, and q the elementary charge. To linearize the I/V characteristic, a large number of $N$ pseudo-resistor elements are connected in series so that $V_{DS}$ does not exceed $U_T$ even for the TIA’s maximum output voltage. Since the pseudo-resistors in A are operated in weak inversion, they exhibit exponential dependencies on absolute temperature and threshold voltage. A specific biasing circuit (B) renders the exponential dependencies on absolute parameters to exponential dependencies on the mismatch between those parameters of the pseudo-resistors and the biasing circuit’s transistors[4]. The large number of elements further averages the transistor’s mismatch and, hence, the resistance value is precise and robust against variations of temperature and process parameters[5]. Furthermore, the biasing circuit facilitates the pseudo-resistors to be floating due to its current source, which provides the floating biasing voltage $V_{bias}$ via the source follower. The entire device of Supplementary Fig. 4 is referred to as multi-element pseudo-resistor (MEPR)[6]. The MEPR’s resistor value can be tuned using an external reference resistor $R_{REF}$. A large tuning range from 1MΩ to 1GΩ has been reported[4]. Moreover, it has been shown in  that the MEPR features a noise floor similar to the Johnson noise of an equivalent ideal ohmic resistor[6]. To minimize the noise, we have implemented the TIA with its maximum gain of 1GΩ.

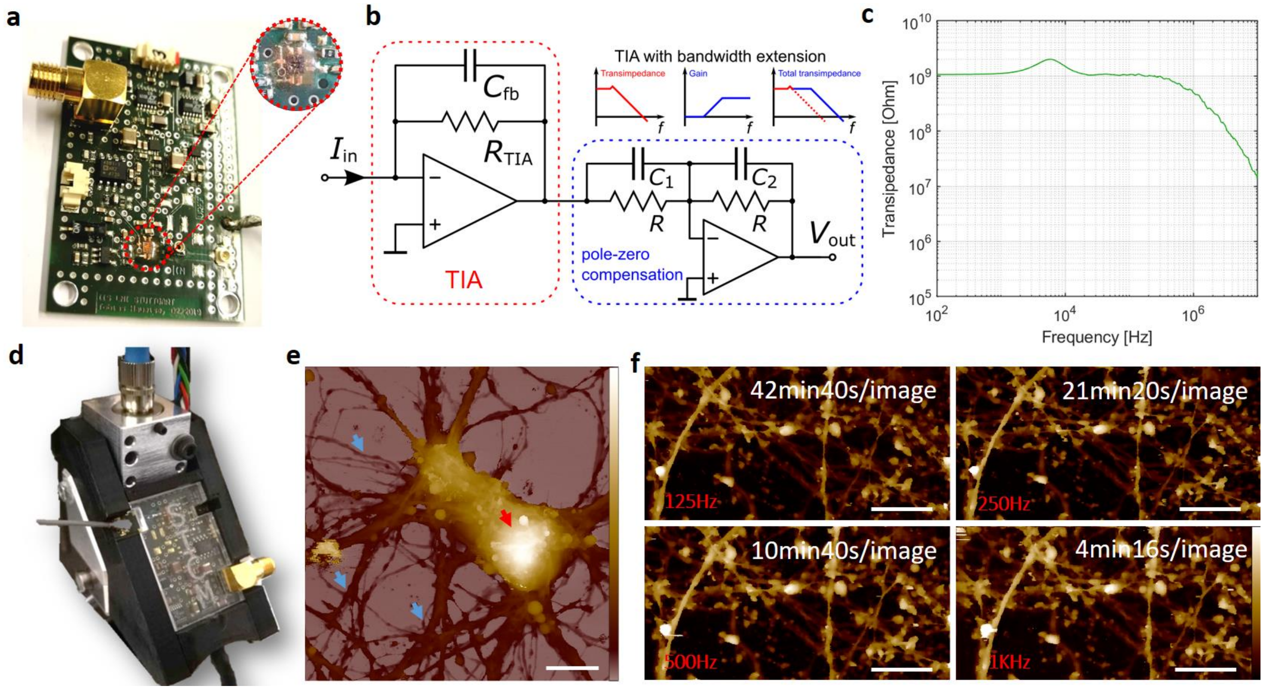


***Supplementary Fig. 5: Custom monolithic bandwidth-extended transimpedance amplifier (BE-TIA) for high-speed current-to-voltage conversion. a****) Printed circuit board and a zoom-in of the BE-TIA.* ***b****) Schematic of the TIA (red) and the succeeding pole-zero compensation circuit for bandwidth extension (blue).* ***c****) Measured transimpedance vs. frequency of the BE-TIA.* ***d****) Integration of the BE-TIA circuit in the SICM setup.* ***e****) To demonstrate the performance of the system, we acquired a 60 µm area image of a single mouse cortical neuron, fixed in 4% PFA in PBS solution. The red arrow shows the cell body and the blue arrows show the intricate network of neurites. Scale bar, 20 µm. Z scale, 0–8 µm.* ***f****) To demonstrate the improvement in frame rate acquisition (512 × 256 pixels) with the integrated BE-TIA circuit, we performed SICM imaging of fixed cortical neurons. This panel shows no loss in the image quality of the neurites for higher hopping rates. Images were acquired at 125 Hz, 250 Hz, 500 Hz, and 1 kHz hopping rate (500 nm hopping height). Scale bar, 10 µm. Z scale, 0–2 µm.*

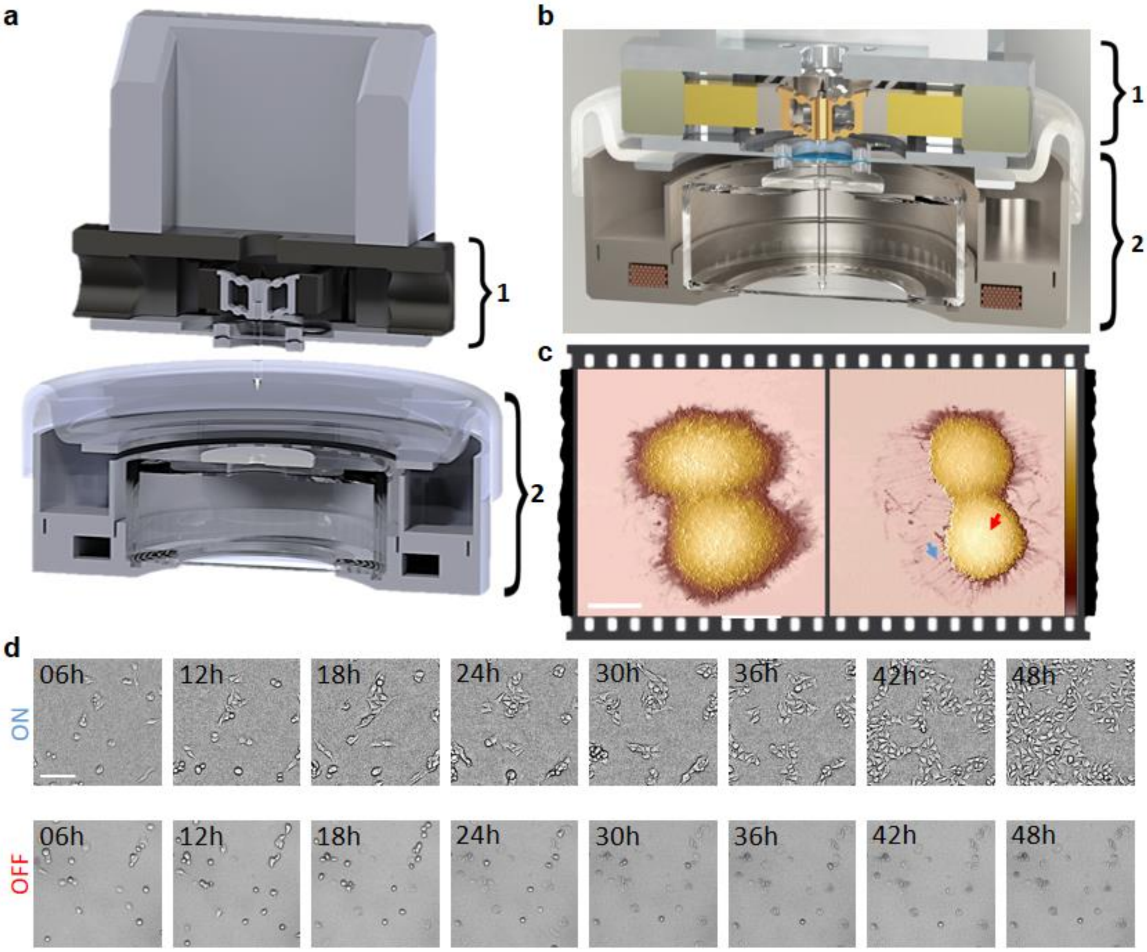


***Supplementary Fig. 6: Time-resolved SICM system composed of a high-performance SICM actuator integrated into an environmental chamber with temperature and $CO_2$ control. a**) 3D rendering of a cross-section of the SICM actuator (1) apart from the miniature incubator (2). **b**) 3D rendering of a cross-section of the enclosed SICM system (1 and 2 together). **c**) Without the appropriate culture environment during the scanning, cells tend to undergo apoptosis within a few hours. This SICM topography image shows HeLa cells detaching from the surface after 6 hours without temperature and $CO_2$ diffusion in cell medium. The red arrow shows the cell body rounding and the blue arrow points to retracted adhesions where the cells were previously attached. Scale bar, 20 µm. Z scale, 0–12 µm. **d**) Upper row: Brightfield microscopy of cell growth under conditions of a controlled atmosphere, demonstrating cell viability and proliferation over 48 hours. Bottom row: Cell growth without environmental control.*

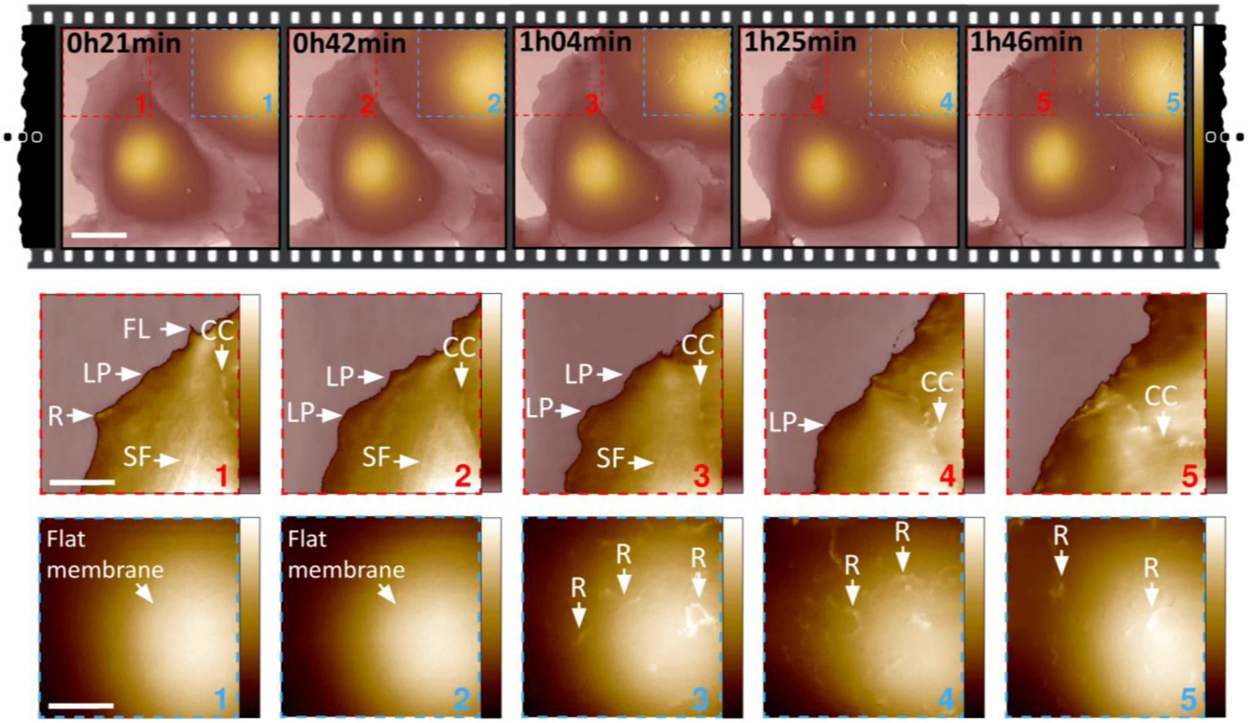


***Supplementary Fig. 7: Time-resolved SICM revealing dynamic protrusions on the apical cell surface.*** *Scale bar, 20 µm. Z scale, 0–8.5 µm. The sequence of zoom-in in red (1-5) shows motility on the cell periphery, with arrows identifying ruffles (R), lamellipodia (LP), filopodia (FL), cell-cell contact (CC), and stress fibers (SF). Scale bar, 10 µm. Z scale, 0–1.2 µm. The sequence of zoom-in in blue (1-5) shows the sudden appearance of dynamic ruffles (R) on the top of the cell membrane. Scale bar, 10 µm. Z scale, 0–5.2 µm.*

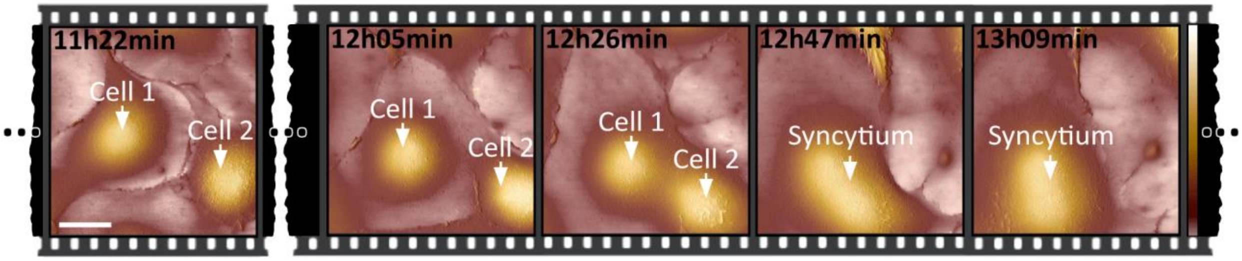


***Supplementary Fig. 8: Time-resolved SICM revealing two cells fusing in syncytium.*** *Arrows point to the top of two cells moving towards each other, fusing in syncytium. Scale bar, 20 µm. z scale, 0–7 µm.*

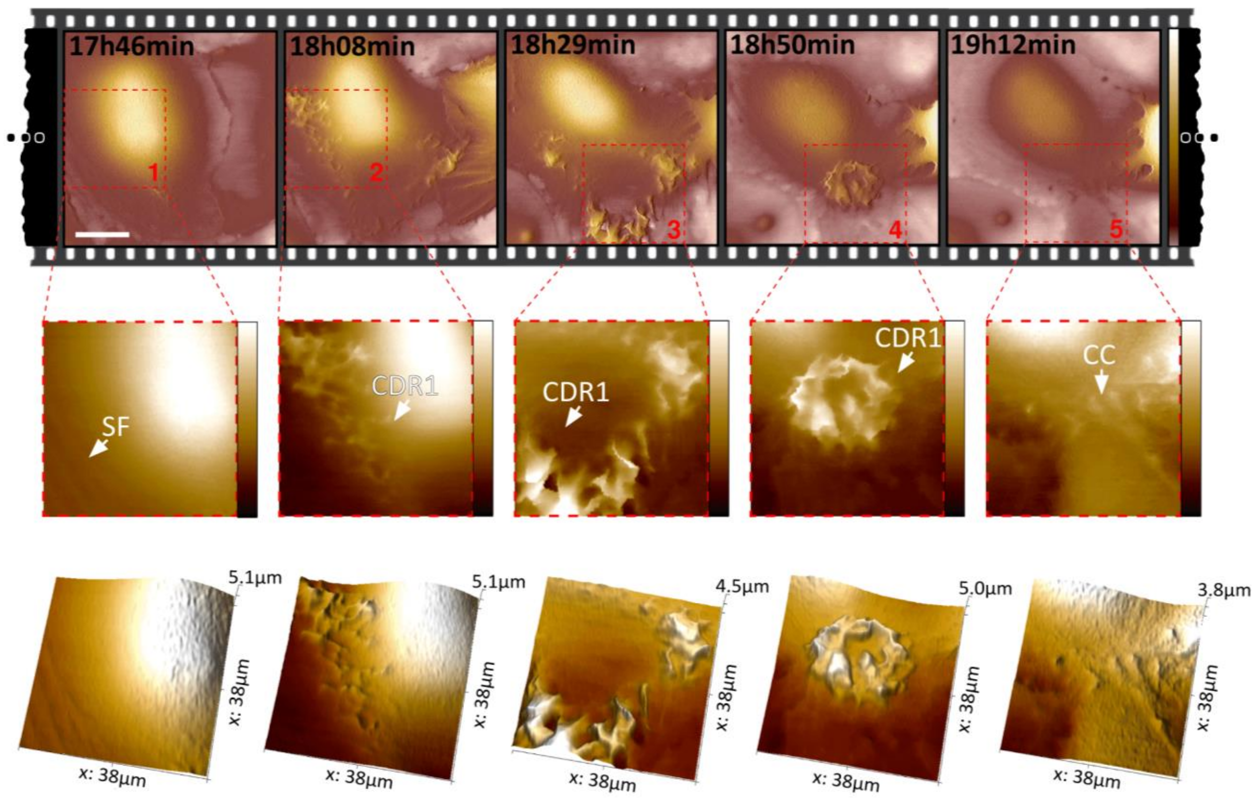


***Supplementary Fig. 9: Time-resolved SICM revealing the appearance and disappearance of circular dorsal ruffles (CDR1).*** *Scale bar, 20 µm. Z scale, 0–7 µm. The sequence of zoom-in in red (1-5) shows the sudden formation of a CDR. Z scale, 0–5 µm, 0–5 µm, 0–4 µm, 0–5 µm, 0–3 µm. Three-dimensional view of the sequence on the bottom.*

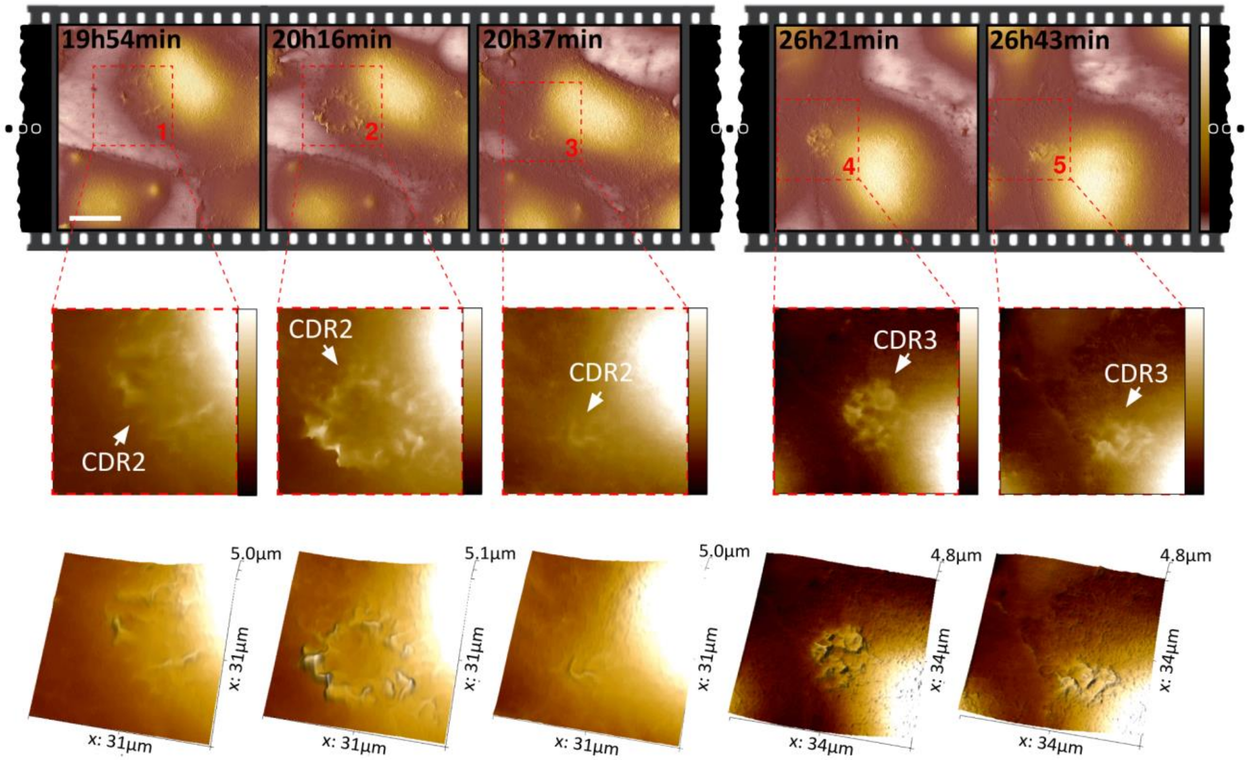


***Supplementary Fig. 10: Time-resolved SICM revealing the appearance and disappearance of two circular dorsal ruffles CDR2, CDR3.*** *Scale bar, 20 µm. Z scale, 0–7 µm. The sequence of zoom-in in red shows the sudden formation of two CDRs, CDR2 (1-3) and CDR3 (4-5). Z scale, -1–4 µm and 0–4 µm for 1-3 and 4-5 respectively. Three-dimensional view of the sequence on the bottom.*

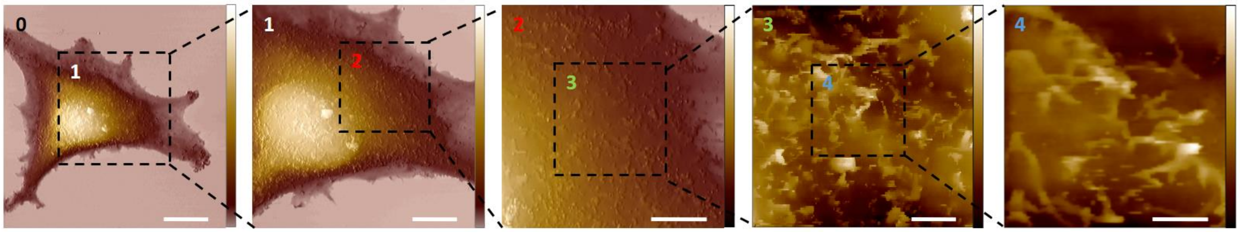


***Supplementary Fig. 11: High dynamic range of XYZ-actuation with SICM high-speed performance.*** *The first image shows the topography of an entire melanoma cell body on an area of 100 µm width. Acquired at 125 Hz hopping rate and 5µm hopping height, 512 × 256 pixels. Z scale, 0-12 µm. Scale bar, 20 µm. 1: Topography image on a smaller area of 50 µm width. Acquired at 200 Hz hopping rate and 3 µm hopping height, 512 × 256 pixels. Scale bar, 10 µm. Z scale, 0-12 µm. 2: Topography image on a smaller area of 20 µm width. Acquired at 285 Hz hopping rate and 2 µm hopping height, 256 × 128 pixels. Scale bar, 5 µm. Z scale, 0-10 µm. 3: Topography image on a smaller area of 10 µm width. Acquired at 500 Hz hopping rate and 1 µm hopping height, 256 × 128 pixels. Scale bar, 2 µm. Z scale, 0-1.5 µm. 4: Topography image on a smaller area at 10 s/image. From a sequence of images acquired at 1 kHz hopping rate and 800 nm hopping height, 100 × 100 pixels. Scale bar, 1 µm. Z scale, 0-1.2 µm. For better visualization, images 0-2 were merged with the slope, and 3-4 were leveled by a mean plane subtraction.*

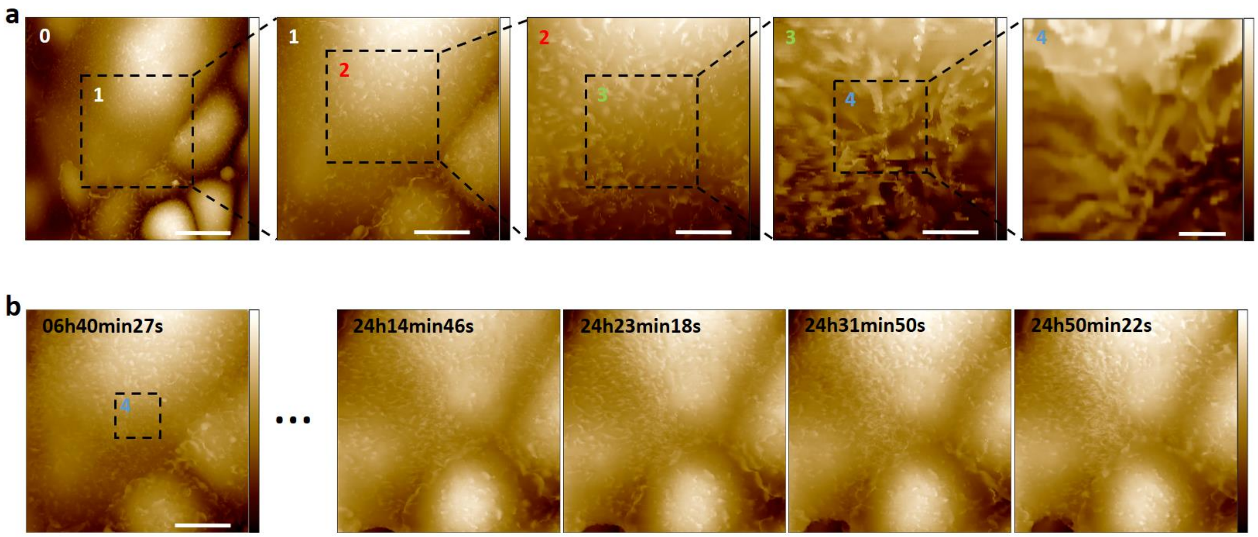


***Supplementary Fig. 12: High dynamic range of XYZ-actuation with SICM high-speed performance and long time-lapse SICM capabilities. a**) The first image shows the topography of kidney cells on an area of 80 µm width. Acquired at 200 Hz hopping rate and 3 µm hopping height, 512 × 256 pixels. Scale bar, 20 µm. Z scale, 0-7.5 µm. 1: Topography image on a smaller area of 40 µm width. Acquired at 250 Hz hopping rate and 2.5 µm hopping height, 512 × 256 pixels. Scale bar, 10 µm. Z scale, 0-7 µm. 2: Topography image on a smaller area of 20 µm width. Acquired at 285 Hz hopping rate and 2 µm hopping height, 256 × 128 pixels. Scale bar, 5 µm. Z scale, 0-4.5 µm. 3: Topography image on a smaller area of 10 µm width. Acquired at 500 Hz hopping rate and 1 µm hopping height, 256 × 128 pixels. Scale bar, 2.5 µm. Z scale, 0-2.5 µm. 4: Topography image on a smaller area. From a sequence of images acquired at 670 Hz hopping rate and 800 nm hopping height, 128 × 64 pixels. Scale bar, 1 µm. Z scale, 0-2 µm. **b**) Zoom-out scanning from a4 followed by a 24 hours time-lapse sequence. Acquired at 250 Hz hopping rate and 2.5 µm hopping height, 512×256 pixels. Scale bar, 10 µm. Z scale, 0-7 µm for the frame at 06h40min27s and 0-5.5 µm for the four frames after 24 hours.*

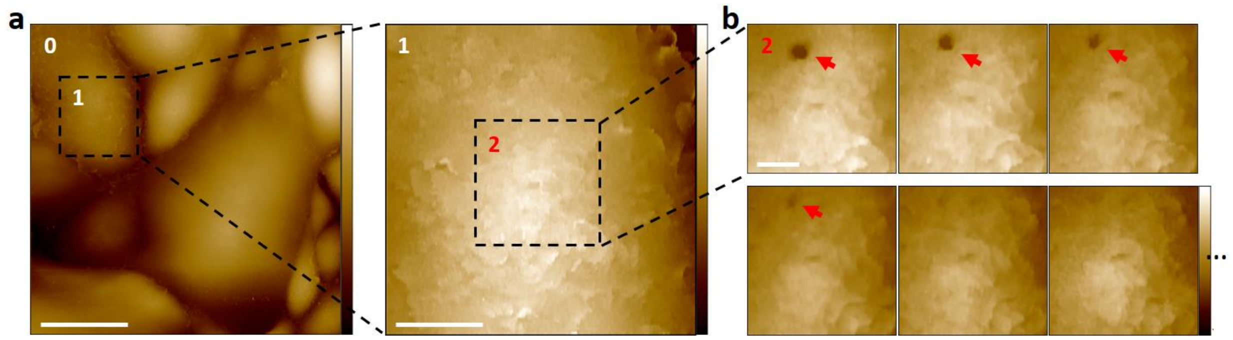


***Supplementary Fig. 13: Increased temporal resolution on detected events of interest. a****) The first image shows the topography of kidney cells on a large area. Acquired at 125 Hz hopping rate and 5 µm hopping height, 512 × 256 pixels. Scale bar, 16 µm. Z scale, 0-10 µm. 1: Topography image on a smaller area, marked in 0. Acquired at 250 Hz hopping rate and 2 µm hopping height, 256 × 128 pixels. Scale bar, 4 µm. Z scale, 0-5 µm.* ***b****) By decreasing the scanning area and focusing on detected events of interest (area marked in 1), we can increase the temporal resolution. Processes such as endocytosis can be visualized on kidney cells (Red arrow). Acquired at 285 Hz hopping rate and 2 µm hopping height, 128 × 64 pixels. Scale bar, 2 µm. Z scale, 0-3 µm.*

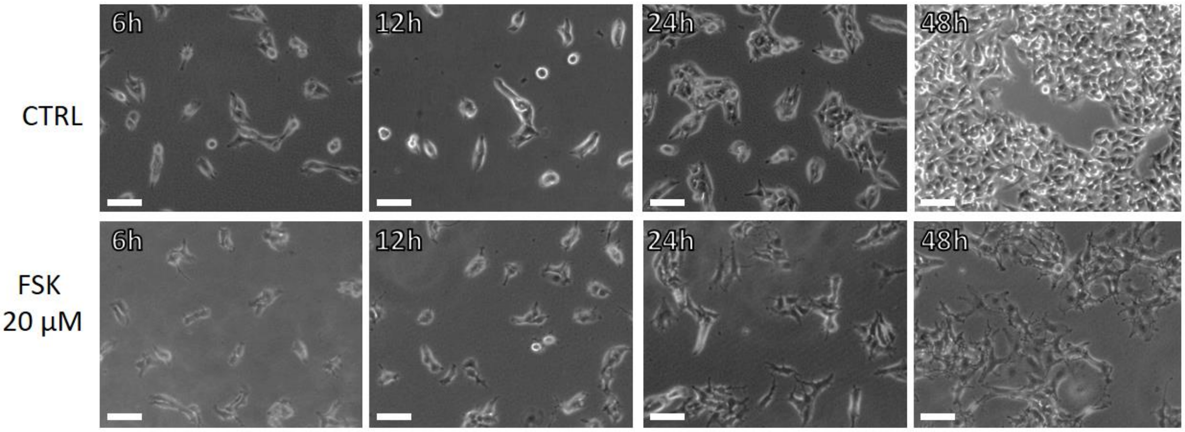


***Supplementary Fig. 14: Effect of Forskolin (FSK) on melanoma cells.*** *50’000 cells were seeded on a glass-bottom petri-dish followed by an attachment period of 6 hours. Then cells were treated with 20 µM FSK (Bottom panel) and compared with the control (Top panel). Image show cells at 6, 12, 24, and 48 hours after cell seeding imaged with a phase-contrast microscope. Scale bar, 200 µm.*

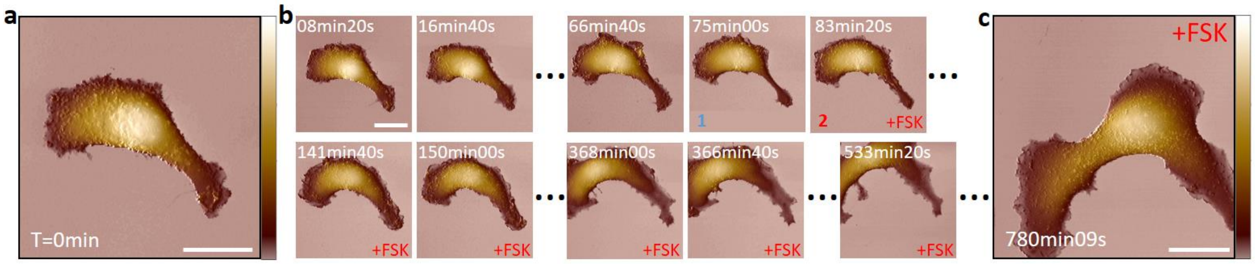


***Supplementary Fig. 15: Time-resolved SICM imaging sequence showing dendrites outgrowth overtime by a melanoma cell treated with FSK. a****) The first image of the sequence before FSK treatment. Scale bar, 10 µm. Z scale, 0–9 µm.* ***b****) Several time points of the sequence over 780 minutes showing the effect of 20 µM FSK (final concentration in the medium) on a single dendrite (1) before and (2) after treatment.* ***c****) The last image of the sequence after 780 minutes highlighting morphological changes induced by FSK treatment, characterized by an increase in the width of the dendrite. Acquired at 125 Hz hopping rate and 6µm hopping height, 300 × 200 pixels. Scale bar, 10 µm. Z scale, 0–9 µm.*

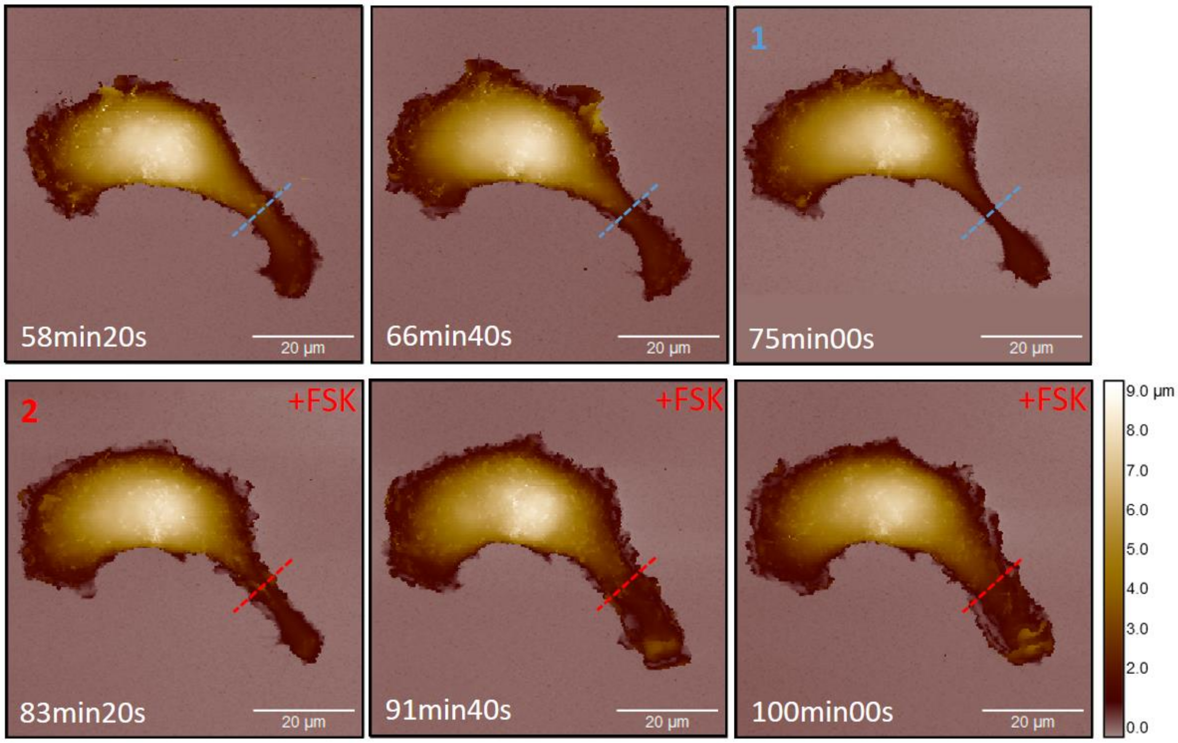


***Supplementary Fig. 16: Topography profiles of dendrites outgrowth on a melanoma cell protrusions after the indicated duration of FSK treatment.*** *Height profile of the dendrite (Fig. 4e) over a time sequence. 1 shows a cell dendrite before treatment and 2 after adding 20 µM FSK.*

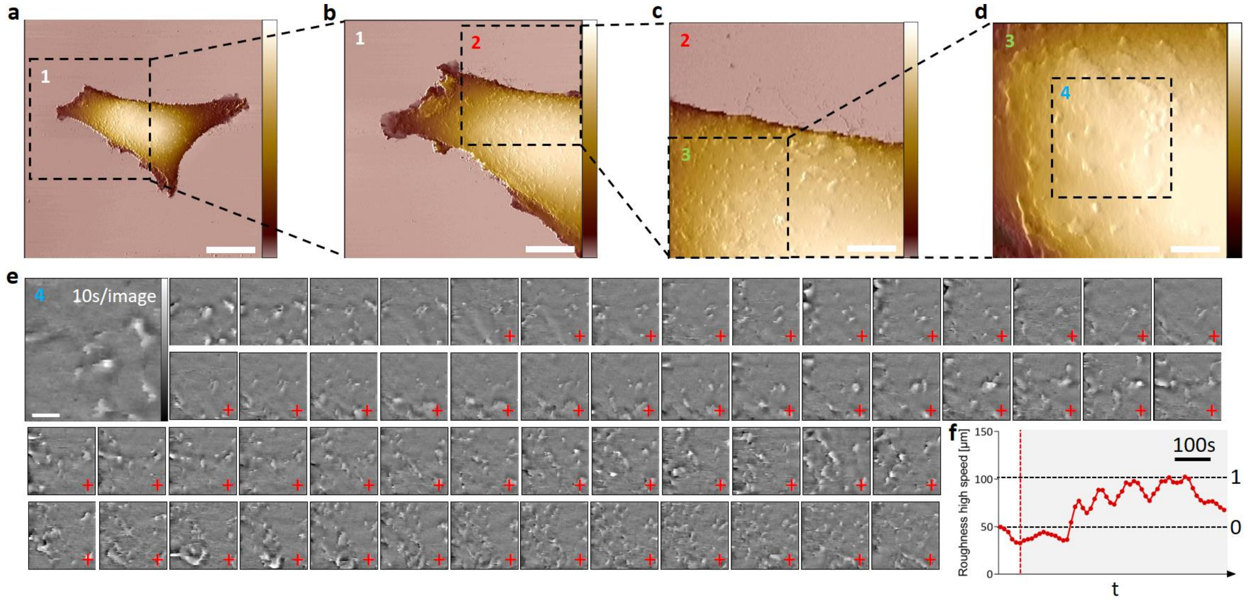


***Supplementary Fig. 17: Real-time effect of FSK on a melanoma cell membrane with fast SICM image acquisition.*** ***a****) Topography image of a melanoma cell on an area of 100 µm. Acquired at 125 Hz hopping rate and 5 µm hopping height, 512 × 256 pixels. Scale bar, 20 µm. Z scale, 0-11 µm.* ***b****) Topography image of the area 1 marked in a. Acquired at 200 Hz hopping rate and 3 µm hopping height, 512 × 256 pixels. Scale bar, 10 µm. Z scale, 0-9 µm.* ***c****) Topography image of the area 2 marked in b. Acquired at 285 Hz hopping rate and 2 µm hopping height, 256 × 128 pixels. Scale bar, 5 µm. Z scale, 0-8 µm.* ***d****) Topography image of the area 3 marked in c. Acquired at 500 Hz hopping rate and 1 µm hopping height, 256 × 128 pixels. Scale bar, 2.5 µm. Z scale, 0-6 µm.* ***e****) Real-time effect of FSK (+) on the melanoma cell membrane of the area 4 marked in d at 10 s/image. From a sequence of images acquired at 1 kHz hopping rate and 500 nm hopping height, 100 × 100 pixels. Scale bar, 1 µm. Z scale, -300 to +300 nm. For better visualization and quantification, the images were flattened and fixed to median zero, followed by a 2-pixel conservative denoise filter.* ***f****) Plot of the surface roughness as a measure to quantify protrusion activity on the cell membrane over time after adding 20 µM FSK (red dashed line). (0) Max roughness before FSK treatment at the beginning as a baseline. (1) Maximal roughness level after FSK treatment. Roughness is defined as the root mean square of height irregularities, computed from 2nd central moment of data values.*

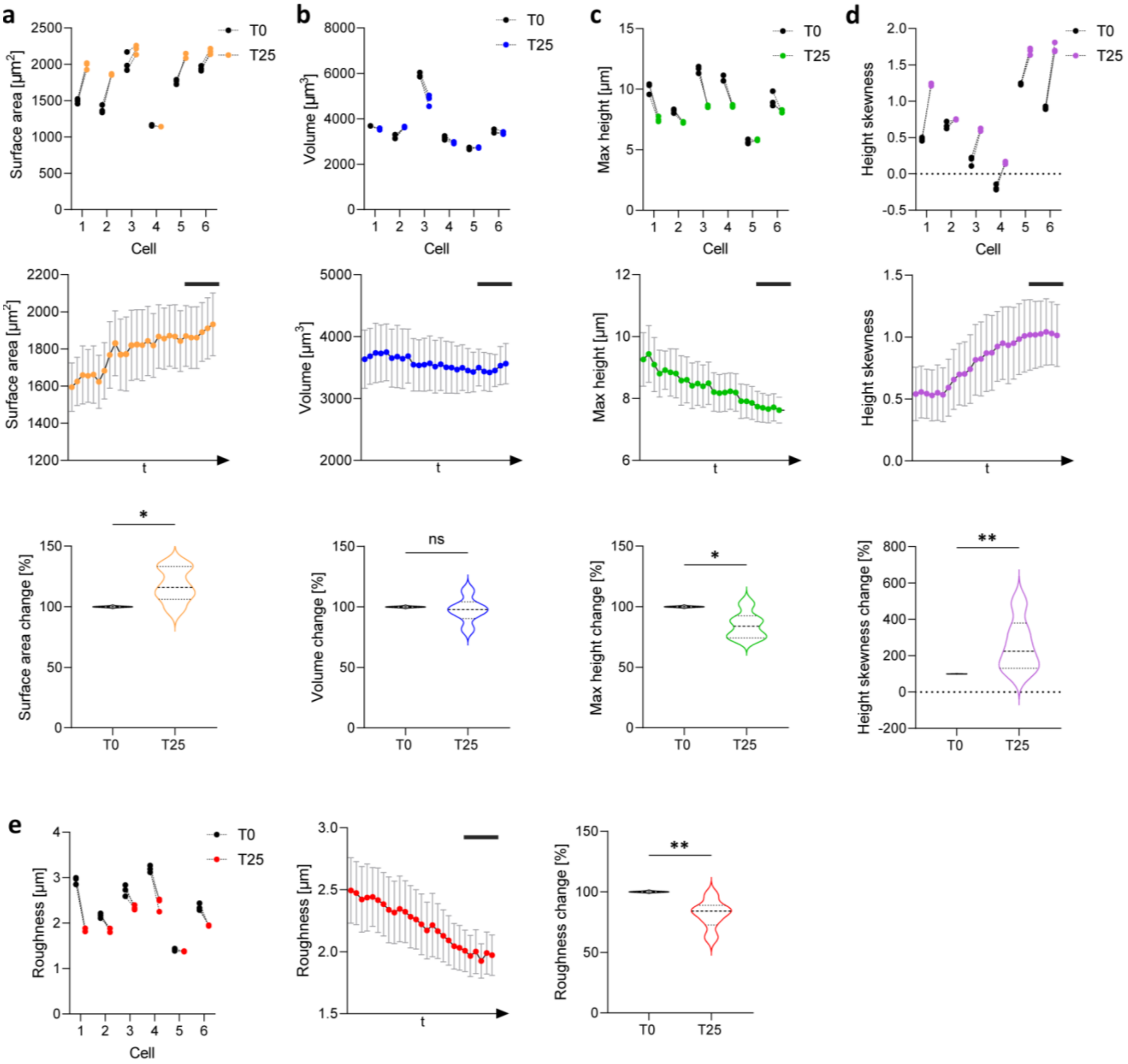


***Supplementary Fig. 18: Tracking morphological changes on melanoma cell treated with FSK with time-resolved SICM. a**) Change in surface area; **b**) volume change; **c**) max height change; **d**) Height skewness change; and **e**) roughness change. Each parameter was measured in 6 different cells. First graph shows measured value in the first three frames of scanning (T0), related to the values measured at frames 25-27 of the same cell (T25). Graph below shows the change in the measurement over time. Violin plots represent percentage of the parameter change at T25 relative to T0. Error bars represent SEM, n=6. *P < 0.05, **P < 0.01. Data compared using two-tailed Mann-Whitney test. Scale bar 50 minutes. The surface area is computed by triangulation of neighboring pixels and the volume is calculated as the integral of the surface height over the covered area. Height skewness is computed from the third central moment of data values.*

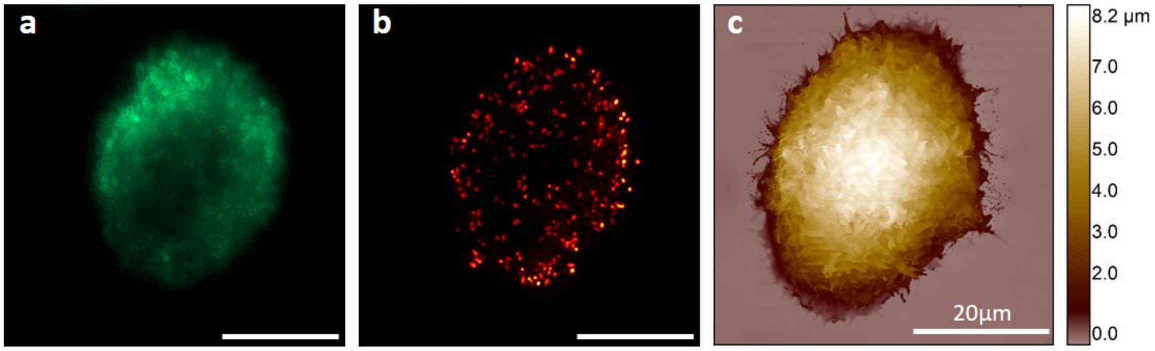


***Supplementary Fig. 19: Fluorescence image of the bacterium affinity (E. coli) to the host mammalian cell membrane (HeLa), correlated with SICM topography. a****) cd80 based GFP display in HeLa cells for direct visualization of the ligand (GFP) sequestered by VHH-intimin expressed by E. coli K12. Scale bar, 10 µm.* ***b****) Visualization of E. coli K12 expressing mScarlet in the cytosol. Scale bar, 10 µm.* ***c****) Three-dimensional representation of the live cell membrane surface with SICM. Acquired at 125 Hz hopping rate and 5 µm hopping height, 512 × 512 pixels.*

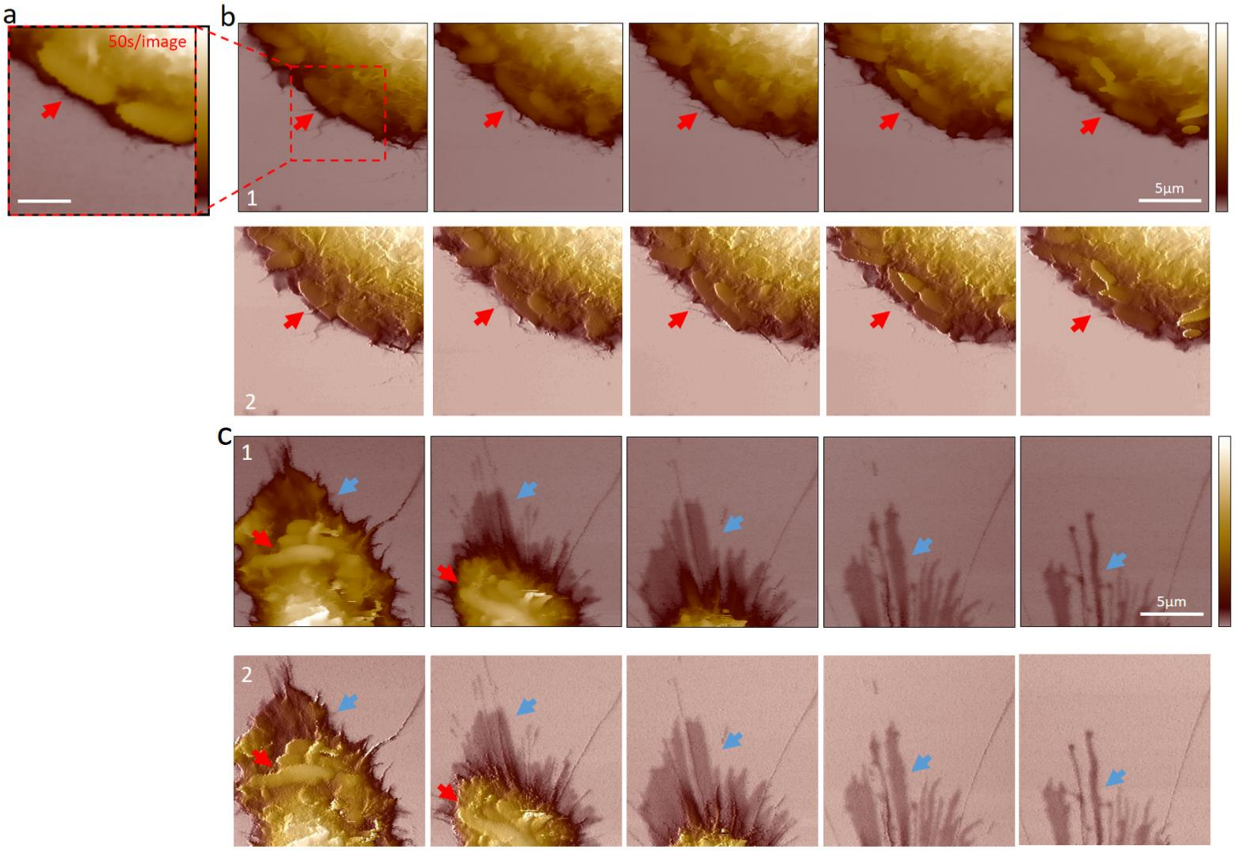


***Supplementary Fig. 20: Time-resolved SICM showing a HeLa cell being infected by E. coli  bacteria. a****)* *Topography image resolving E. coli dividing on the host HeLa cell membrane on a small area of 7.5 µm. Acquired at 250 Hz hopping rate and 2 µm hopping height, 100 × 100 pixels. Scale bar, 2 µm.  Z scale, 0-3.2 µm.* ***b****) Topography of E. coli on a 15 µm area adhering and proliferating on the host cell membrane periphery (1). Red arrows point to the bacteria dividing on the membrane. Acquired at 125 Hz hopping rate and 4 µm hopping height, 256×256 pixels. Scale bar, 5 µm. Z scale, 0–6 µm.* ***c****) Topography of E. coli adhered on the host cell membrane dendrite (1). Red arrows point to bacteria on the membrane and blue arrows point to the dendrite retracting. Acquired at 125 Hz hopping rate and 4 µm hopping height, 256×256 pixels. For better visualization, the topography channel was merged with the slope channel to enhance the surface edges (2). Scale bar, 5 µm. Z scale, 0–3 µm.*

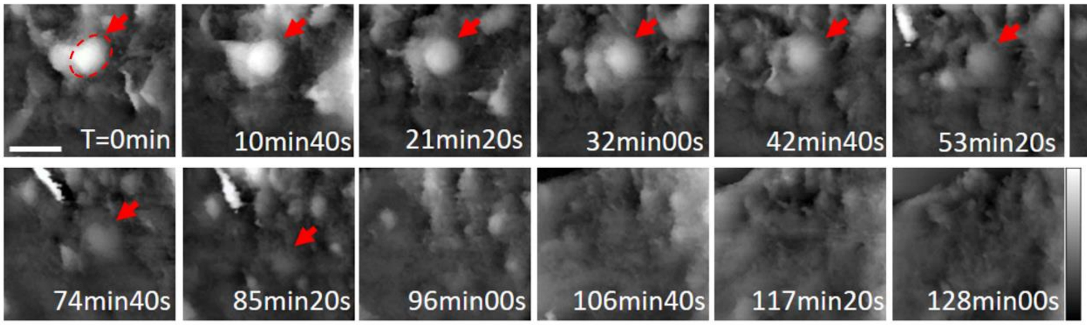


***Supplementary Fig. 21: Time-resolved SICM showing an E. coli being internalized by HeLa cell membrane over time.*** *Arrows point to a bacterium being internalized over 85 minutes. Zoom-in cut on an area from the long time-lapse shown in Fig. 5e. Acquired at 125Hz hopping rate and 4 µm hopping height, 256×256 pixels. For better visualization, the membrane surface was flattened. Scale bar, 2 µm. Z scale, 0-1.1 µm.*

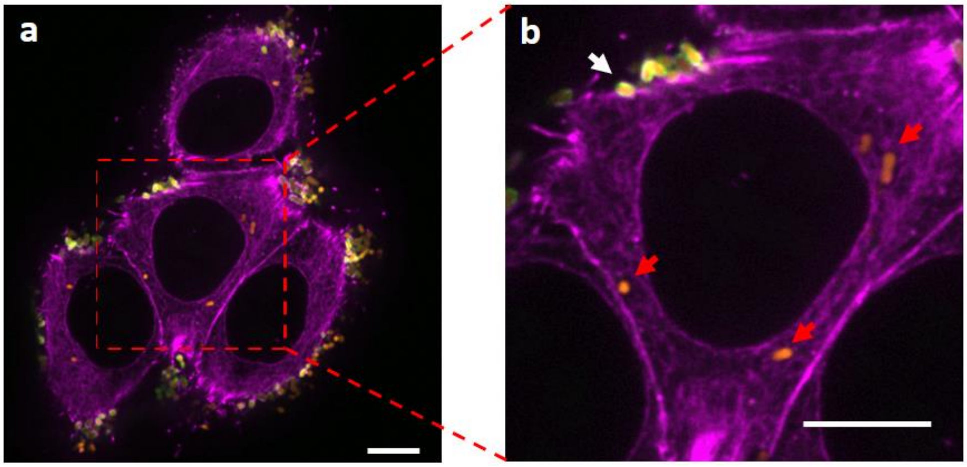


***Supplementary Fig. 22: Fluorescence image of an E. coli internalized by HeLa cell membrane. a****)* *Infected cell showing bacteria adhered on the membrane and locally accumulating GFP signal (cd80 based GFP display). Cells were fixed in 4% paraformaldehyde for 20 minutes, permeabilized with 0.1% Triton X-100 for 5 minutes, and washed twice with PBS. Phalloidin-Atto 655 (Sigma) was used to stain actin at 500 nM for 15 minutes (purple). Data collected with a spinning disk confocal microscope and 100x oil immersion objective. Scale bar, 10 µm.* ***b****) Zoom in on an area showing bacteria that were internalized and lost the GFP signal (red arrows), potentially due to lysosomal pH. The white arrow shows bacteria that were not internalized and kept the GFP signal. Scale bar, 10 µm.*

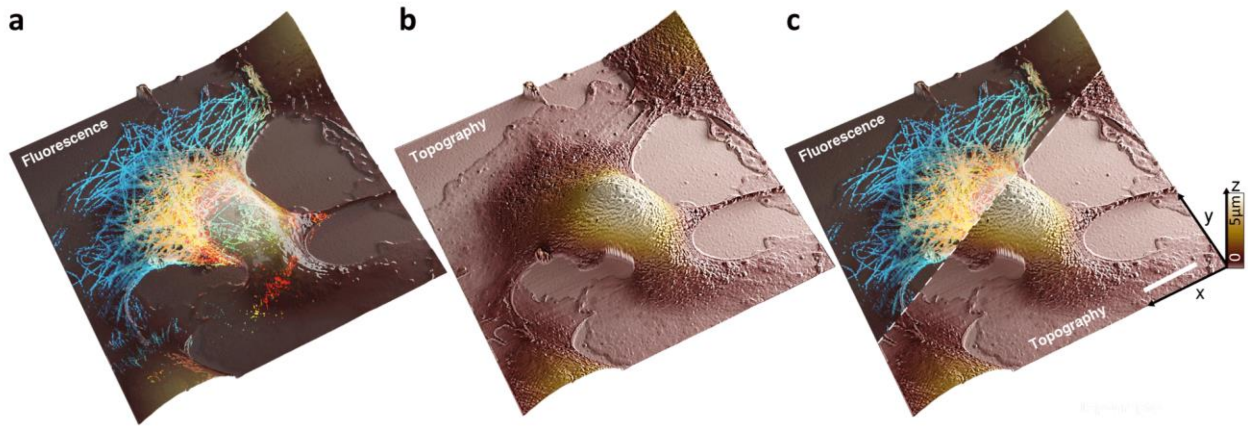


***Supplementary Fig. 23: Correlative fluorescence and scanning ion conductance microscopy. a****) Fluorescence image of microtubules.* ***b****) SICM topography image of the cell membrane.* ***c****) Correlative fluorescence/topography image of a mammalian cell (COS7). Scale bar, 10 µm.*
